# An EEG-fMRI Jointly Constrained Digital Twin Brain and Its Application in Alzheimer’s Disease

**DOI:** 10.64898/2026.04.10.717365

**Authors:** Yue Xiong, Daqing Guo, Yuanhang Xu, Yutao Chen, Rongxin Zhang, Yiqian Luo, Feiyan Wang, Xin Zeng, Yi Guo, Dezhong Yao

## Abstract

Current digital twin brain (DTB) models are typically optimized using single-modality functional neuroimaging data, which restricts their ability to simulate brain dynamics across multiple spatiotemporal scales. Here, we show that a two-stage DTB (TS-DTB) modeling framework jointly constrained by functional magnetic resonance imaging (fMRI) and electroencephalography (EEG) data. Validation in both the healthy young and Alzheimer’s disease (AD) cohorts demonstrates that the TS-DTB model simultaneously captures subject-specific features of multi-scale brain dynamics. In particular, the TS-DTB models of AD patients successfully recapitulate spectral-temporal signatures of cognitive decline, suggesting a model-inferred association between these deficits and simulated excitation-inhibition (E-I) imbalances. By simulating responses to repetitive transcranial magnetic stimulation (rTMS), we further reveal that acute EEG normalization in AD patients is computationally associated with putative E-I rebalancing via model-inferred synaptic reconfiguration and background suppression. Overall, these findings underscore the potential of the TS-DTB to advance the mechanistic understanding of the brain and inform personalized digital therapeutics.

## Introduction

The digital twin brain (DTB) has emerged as a transformative modeling framework for investigating brain dynamics, offering a distinct research perspective on brain function, cognitive behaviors, disease mechanisms, and neuromodulation strategies^1–3^. By coupling structural connectivity with biologically grounded dynamical descriptions, the DTB model reproduces rich macroscopic nonlinear dynamics, including oscillations, synchronization, bifurcations, and critical transitions^4,5^. Driven by the clinical demand for precision medicine, recent advances have increasingly prioritized integrating multiscale architectures and multimodal constraints, evolving from static network fitting toward the reconstruction of dynamic brain states^6–8^. To support this evolution, researchers have systematically incorporated biological priors, such as intrinsic hierarchy and structure–function relationships, into these models to constrain the optimization of high-dimensional parameter spaces^8–11^. The ultimate goal is to elucidate the underlying mechanisms of brain function and disease, predict brain responses, and design personalized, model-driven interventions^12–14^.

Despite these advances, effectively integrating the complementary insights of multimodal neuroimaging remains a fundamental challenge^15^. The fMRI delineates the hemodynamic network topology with superior spatial resolution, whereas EEG reproduces the rapid spatiotemporal dynamics of neuronal activity^16,17^. However, existing DTB modeling strategies often rely on single-modality information constraints, forcing a compromise between spatial topology and temporal fidelity. For instance, models optimized solely for fMRI functional connectivity frequently fail to generate realistic fast oscillations^9,10,18^. Conversely, those tuned exclusively for EEG or magnetoencephalography (MEG) spectral-temporal features often compromise the underlying spatial network architecture^6,19^. This spatiotemporal disconnect creates a fundamental blind spot, preventing current models from reproducing the subtle interplay between network and fast oscillatory dynamics^20^. Resolving this multi-scale interaction is a critical prerequisite for realizing the clinical potential of DTB, particularly for deciphering complex pathologies such as Alzheimer’s disease (AD)^21,22^. Consequently, while progress has been made in multimodal integration^6,9,23,24^, achieving a unified modeling framework that reproduces multi-scale spatiotemporal dynamics remains a critical bottleneck.

Overcoming this bottleneck requires the rational parameterization of spatially heterogeneous intra-regional dynamics and inter-regional coupling. The human brain exhibits profound regional heterogeneity in synaptic architecture and local dynamics, a complexity often oversimplified by homogeneous parameter assumptions in early DTB models^19,23,25^. Capturing this heterogeneity is indispensable for studying neurodegenerative diseases, which are characterized by spatially specific disruptions in local excitation-inhibition (E-I) balance^11,26,27^. However, region-specific parameter estimates that deviate from biological realism inevitably lead to severe overfitting and computational intractability. To break this impasse, a robust strategy must biologically constrain local circuit heterogeneity using spatiotemporal priors, while simultaneously employing precise forward modeling to project simulated source activities onto observable hemodynamics and sensor-space EEG signals^28,29^. Such an approach is essential for developing biologically interpretable models that can serve as testbeds for brain research and the theoretical reference for clinical diagnosis and treatment.

To address these challenges, we developed a two-stage DTB (TS-DTB) modeling framework jointly constrained by fMRI and EEG data. We define this digital twin brain as an individualized, multimodal data-constrained computational replica that retrospectively reconstructs a subject’s baseline spatiotemporal brain dynamics and forward-simulates macroscale neurophysiological responses to localized virtual interventions. We conducted systematic validations across healthy young and AD cohorts to evaluate this framework. In the healthy cohort, the individualized TS-DTB models demonstrated superiority over conventional single-modality and homogeneous models by simultaneously reproducing subject-specific, multi-scale brain dynamics. Extending this validation to a pathological regime, the TS-DTB models of AD patients successfully recapitulated the macroscopic spectral-temporal signatures of cognitive decline, suggesting a model-inferred association between these deficits and regional E-I imbalances. Building upon this pathological insight, we further deployed the TS-DTB models as an in-silico clinical testbed to simulate responses to repetitive transcranial magnetic stimulation (rTMS). The simulation revealed that acute EEG normalization in AD is computationally associated with putative E-I rebalancing via model-inferred synaptic reconfiguration and background suppression. Ultimately, the TS-DTB establishes a mechanistically interpretable bridge linking macroscopic neuroimaging to microcircuit dynamics.

## Results

### System overview

The foundation of the TS-DTB modeling framework is a biophysically grounded generative model designed to simultaneously simulate macroscopic neuroimaging and fast electrophysiological signals (Fig. 1a). Whole-brain dynamics were simulated using connectivity-coupled Wilson-Cowan oscillators. These local neural populations generated excitatory firing rates, serving as a biophysical proxy for local field potentials^22^, which were subsequently transformed into observable signals via modality-specific forward models. Specifically, the simulated neural activity was projected into BOLD signals using the Balloon-Windkessel model. Concurrently, to capture sensor-space EEG signatures, the source-level electrical activity was mapped to the scalp using individualized leadfields derived from subject-specific EEG forward models, thereby ensuring spatial fidelity.

**Fig. 1.**
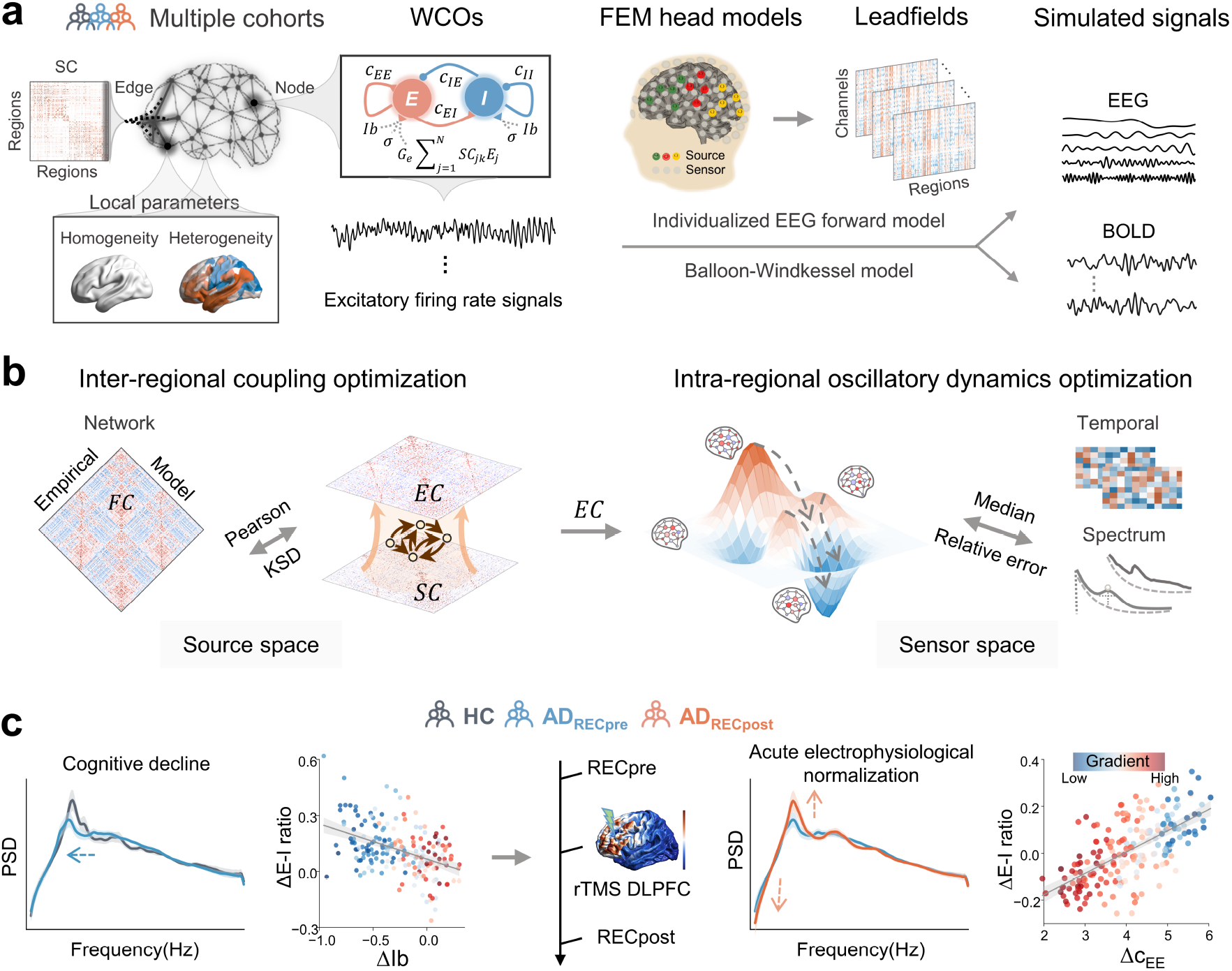
Overview of the TS-DTB modeling framework. **(a)** Construction of individualized DTB models. Whole-brain dynamics were simulated via connectivity-coupled Wilson‒Cowan oscillators (WCOs). Local circuit parameterization incorporated both homogeneous (control) and heterogeneous schemes, with the latter derived from fMRI-based functional gradients. Simulated local excitatory firing rates were converted into EEG and BOLD signals via the individualized EEG forward model and Balloon‒Windkessel model, respectively. **(b)** Schematic of two-stage parameter optimization strategy. The first stage refines the effective connectivity (EC) matrix using an fMRI-based loss function combining the Pearson correlation and the Kolmogorov‒Smirnov distance (KSD) of functional connectivity (FC). Subsequently, the second stage accomplishes individualized kinetic parameterization via Bayesian optimization constrained by a joint loss function derived from the median relative errors (REs) of EEG temporal and spectral features. **(c)** Validation across healthy control (HC) and Alzheimer’s disease (AD) cohorts. The TS-DTB was applied to investigate the neurophysiological mechanisms associated with cognitive decline (left, HC vs. AD_RECpre_) and acute electrophysiological normalization induced by repetitive transcranial magnetic stimulation (rTMS) (right, AD_RECpre_ vs. AD_RECpost_).

To accurately parameterize this high-dimensional dynamic system, we implemented a two-stage optimization strategy utilizing empirical resting-state multimodal features as benchmarks (Fig. 1b, Tables S1-S3). Recognizing that structural connectivity derived from white matter tractography often imperfectly captures inter-hemispheric communication^8,9^, the first stage refined the source-space effective connectivity (EC) constrained by empirical fMRI functional connectivity (FC)^8^. Building upon this optimized network backbone, the second stage focused on reproducing fast sensor-space dynamics by tuning heterogeneous local kinetic parameters. Crucially, this stage leveraged macroscopic fMRI-based gradients as spatial priors. This approach biologically constrains the distribution of regional synaptic strengths, background inputs, and noise to a low-dimensional manifold governed by global gradient weights^11,26^. Subsequently, we employed Bayesian optimization to identify the optimal configuration of these weights, minimizing the relative error (RE) against a broad suite of individualized empirical EEG spectral-temporal features. By synergizing large-scale network calibration with precise local electrophysiological tuning, the TS-DTB ensures that simulated neural activity authentically reconstructs multiscale brain states.

Guided by this architecture, we systematically evaluated the TS-DTB through validation across healthy and AD cohorts (Fig. 1c). In the healthy cohort, we established its methodological robustness by demonstrating that the individualized TS-DTB models exhibit superiority over conventional single-modality models, homogeneous and null baselines, as well as models lacking individualized EEG forward mapping, in accurately reproducing subject-specific spatiotemporal dynamics. Transitioning to the AD cohort, we utilized these models to map macroscopic spectral-temporal abnormalities to underlying disruptions in local circuit parameters. Finally, we deployed the TS-DTB models as an in-silico testbed to simulate rTMS interventions, aiming to track the spatiotemporal responses and investigate the model-inferred parametric shifts associated with acute electrophysiological normalization.

### Reconstructing spatiotemporal dynamics with multimodal constraints

As outlined in the system overview, a fundamental premise of TS-DTB is its ability to integrate EEG–fMRI information for constraining parameter optimization. To evaluate the effectiveness of the multimodal constraint strategy and the chosen optimization order, we compared the TS-DTB models against the unconstrained baseline (noOpt), the EEG-specific (EEGOpt), the fMRI-specific (fMRIOpt) models, as well as an alternative reversed-order scheme (rev-He_TS-DTB_). For instance, while the fMRIOpt models achieved the lowest fMRI loss (defined in Eq. 10, Mean±SE=0.233±0.014), they exhibited substantial deviations in EEG spectral reconstruction (RE = 0.766 ± 0.059). Conversely, the EEGOpt models significantly reduced spectral errors but compromised the underlying spatial connectivity structure, resulting in a high fMRI loss comparable to the unoptimized baseline. Notably, while the second-stage EEG optimization in our TS-DTB model partially compromises the pristine fMRI fit obtained in the first stage, it outperforms the reversed sequential order, which failed to adequately minimize the EEG spectral-temporal losses. The TS-DTB model reconciles this domain-specific trade-off, maintaining a robust fMRI fit (0.933±0.035) while simultaneously achieving the lowest RE for EEG temporal (0.532 ± 0.034) and spectral (0.199 ± 0.031) features. A detailed breakdown of the fMRI loss sub-components and the convergence curves for both optimization stages are provided in Fig. S2. These comparisons confirm that the sequential fMRI-first/EEG-second strategy achieves a global trade-off across multiscale spatiotemporal features, balancing macroscopic structural constraints with local electrophysiological fidelity.

The necessity of regional heterogeneity was further highlighted by the reconstruction of the relative power spectral density (PSD). Models incorporating parameterized heterogeneity (EEGOpt and He_TS-DTB_) successfully aligned with the dominant alpha peak observed in the resting-state brain. In contrast, structurally homogeneous models produced unrealistic broadband oscillatory patterns with high *1/f* noise floors (Fig. 2b). Systematic evaluation across diverse time-series and spectral feature sets (Tables S2-S3) confirmed that the He_TS-DTB_ models consistently minimized deviations specifically for PSD and temporal autocorrelation metrics, effectively capturing the complex, non-linear dynamical signatures of the brain (Fig. 2c).

**Fig. 2.**
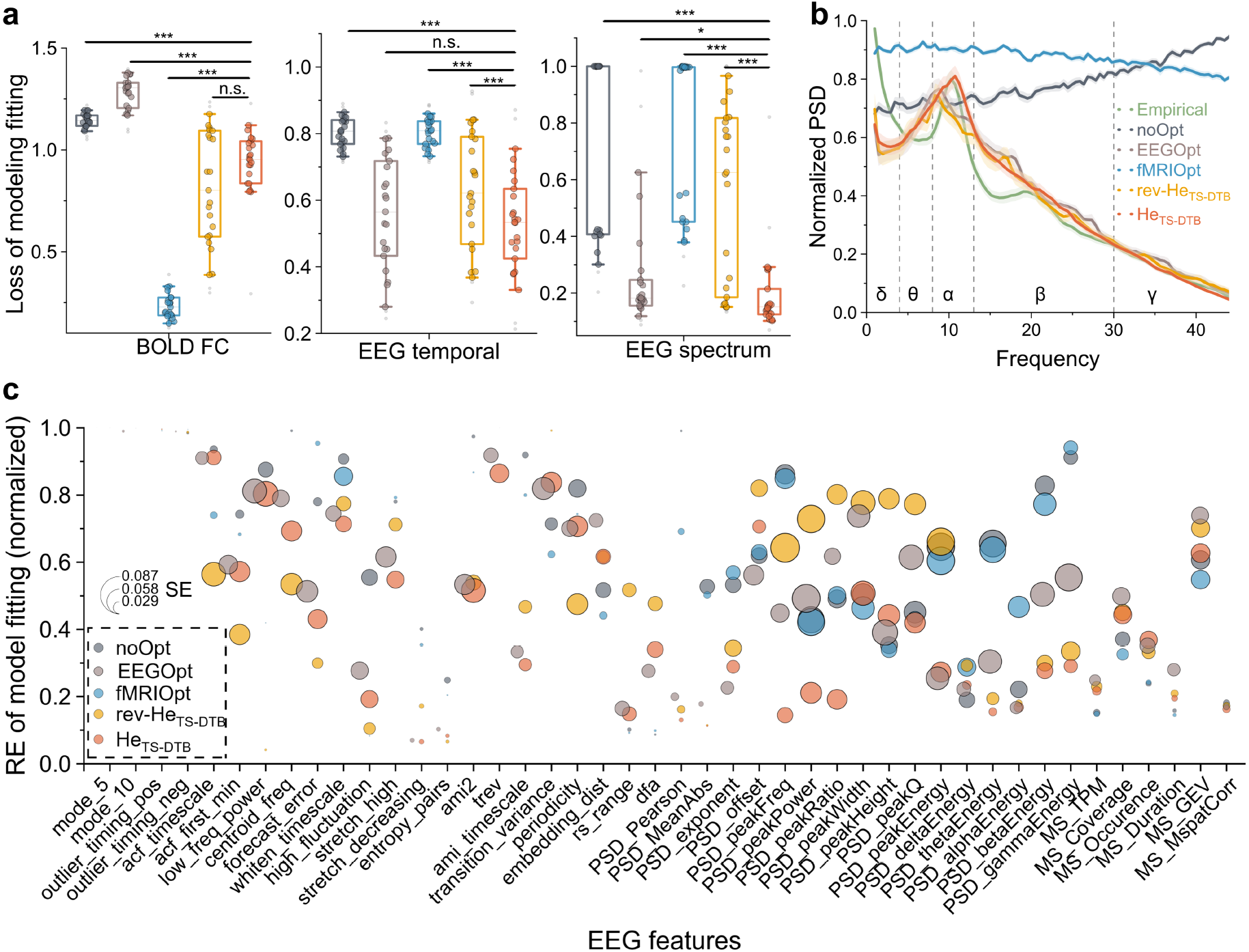
Performance comparison of TS-DTB models against single-modality and alternative optimization schemes. Quantitative evaluation across five parameterization schemes: the proposed models (He_TS-DTB_, i.e., fMRI-first/EEG-second), the reversed-order optimization models (rev-He_TS-DTB_, i.e., EEG-first/fMRI-second), the unconstrained baseline (noOpt), the EEG-constrained models optimizing only heterogeneous local parameters (EEGOpt), and the fMRI-constrained models optimizing only EC coupling (fMRIOpt). **(a)** Global fitting performance. Comparison of the loss function values for fMRI BOLD FC and the median REs for EEG temporal and spectral features. Boxplots indicate the median (center line) and interquartile range (IQR, box limits), whiskers show 90% confidence intervals, and dots denote biologically independent individuals (n = 25). Significance was assessed via two-sided paired t-tests for BOLD FC and EEG temporal, and two-sided Wilcoxon signed-rank tests for EEG spectral (unadjusted for multiple comparisons; each alternative scheme was independently compared only against the proposed He_TS-DTB_ model; \**p* < 0.05, \*\**p* < 0.01, \*\*\**p* < 0.001, n.s. indicates not significant). Exact *p* values (from left to right): BOLD FC loss, *p* < 0.0001, *p* < 0.0001, *p* < 0.0001, *p* = 0.05087; EEG temporal RE, *p* < 0.0001, *p* = 0.161, *p* < 0.0001, *p* = 0.00029; EEG spectral RE, *p* < 0.0001, *p* = 0.04512, *p* < 0.0001, *p* < 0.0001. **(b)** Spectral reconstruction. Channel-averaged relative power spectral density (PSD) across five frequency bands. Solid lines represent the group mean and shaded regions represent the standard error (SE) across all 25 subjects. **(c)** Feature-wise REs. Detailed breakdown of RE for catch22 time-series features, PSD, and EEG microstate metrics. Circle centers indicate the mean RE, and diameters are proportional to the SE (n = 25).

To determine whether this performance improvement relies on biological organization rather than mere parameter expansion, we compared the He_TS-DTB_ models against the null models employing homogeneous (Ho) parameters and the spatially randomized gradients (He_Moran_)^30^. The inclusion of authentic functional gradient priors significantly enhanced optimization, with the He_TS-DTB_ models achieving the lowest minimum loss function values (0.699 ± 0.187) and yielding significantly lower spectral errors (0.199 ± 0.154) compared to both null models (Fig. 3a). Crucially, the inability of the He_Moran_ models to consistently outperform the homogeneous underscores that the enhanced fitting of He_TS-DTB_ models is not a trivial consequence of increased parameter degrees of freedom. Instead, accurate reconstruction of brain dynamics fundamentally depends on constraining local circuit heterogeneity with the intrinsic hierarchical priors of the brain. This biologically grounded parameterization enabled the He_TS-DTB_ models to reconstruct fine-grained details of EEG PSD, including center frequency, spectral energy, and aperiodic components (Fig. 3b). Extending this evaluation across a broader feature set confirmed that authentic spatial hierarchical priors are essential for reproducing complex temporal correlations, signal complexity, and microstate transition properties (Fig. 3c).

**Fig. 3.**
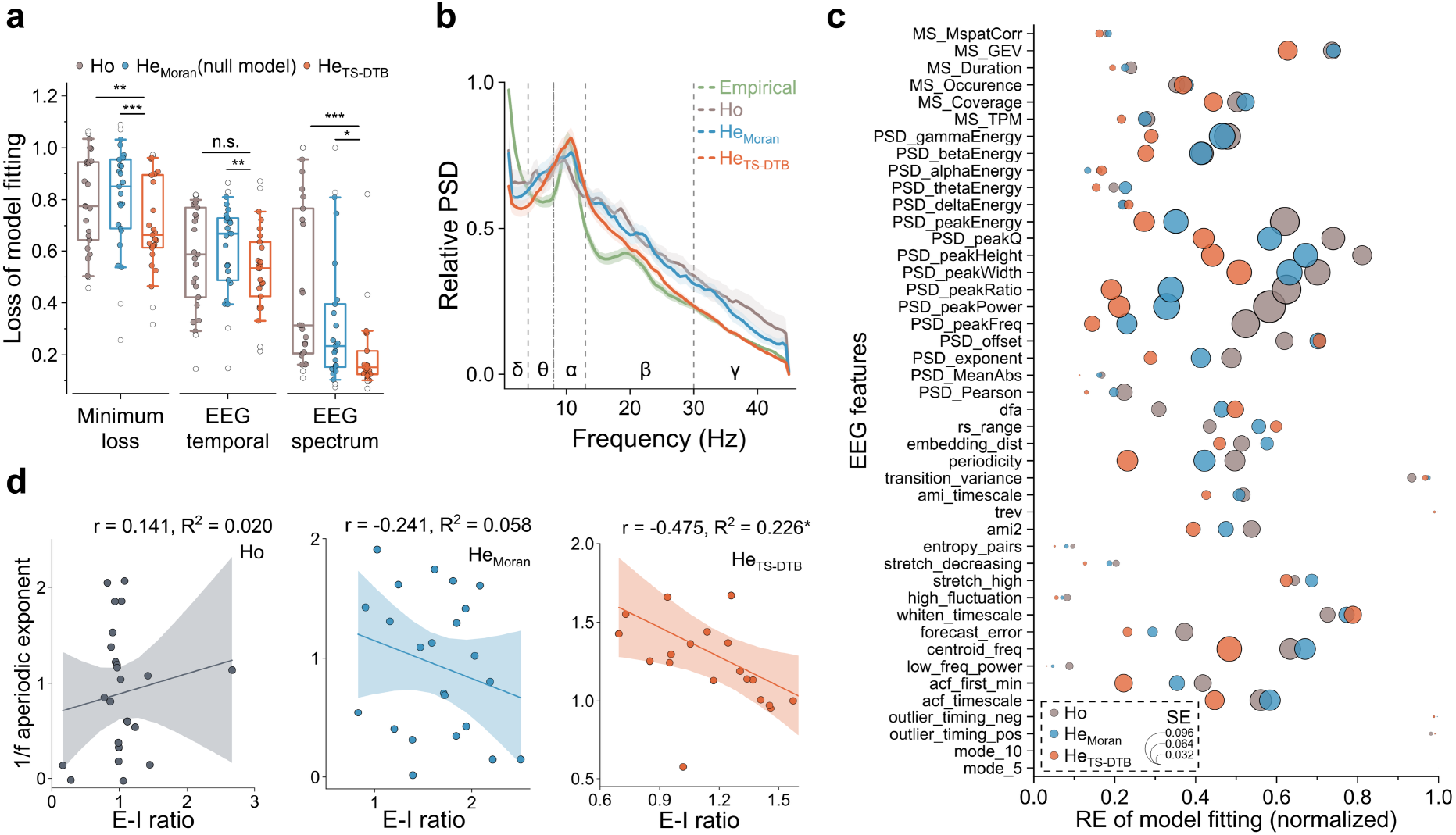
Evaluation of gradient-informed heterogeneity against homogeneous and null models. Model performance under three spatial parameterization schemes, using the identical optimal EC matrix derived from the first stage optimization: the proposed models (He_TS-DTB_), the autocorrelation-preserving spatially randomized null models (He_Moran_), and the homogeneous baseline (Ho). **(a)** Optimization performance. Minimal loss function values and the median RE for EEG features. Boxplots indicate the median (center line) and IQR (box limits), whiskers show 90% confidence intervals, and dots denote biologically independent individuals (n = 25 HC subjects); significance was assessed via two-sided paired t-tests (\*\*\**p* < 0.001). **(b)** Spectral reconstruction. Channel-averaged relative PSD for empirical data and simulations. Solid lines represent the group mean and shaded regions represent the standard error (SE) across subjects (n = 25). **(c)** Feature-wise REs for EEG temporal, spectral, and microstate properties (detailed in Tables S2-S4). Circle centers indicate the mean RE and diameters are proportional to the SE (n = 25). **(d)** Mechanistic validation of E-I balance. Scatter plots illustrating the relationship between the simulated E-I ratio and the aperiodic exponent. The solid line indicates the linear regression fit with 95% confidence intervals (shaded) evaluated via two-sided Pearson correlation across n = 25 biologically independent individuals.

Beyond fitting accuracy, we validated the biophysical plausibility of these models by examining the mechanistic link between the simulated E-I ratio and the *1/f* aperiodic spectral exponent. Empirical and theoretical studies posit that the aperiodic slope reflects the local E-I balance, with stronger inhibition resulting in steeper *1/f* slopes and consequently lower E-I ratios^31,32^. While both the Ho and He_Moran_ models failed to capture this physiological relationship (*p* > 0.05), the He_TS-DTB_ models demonstrated a significant negative correlation (*r*=−0.475, *p* < 0.05) consistent with empirical expectations (Fig. 3d). Further analysis revealed that these local E-I characteristics were primarily driven by the interplay between background input and inhibitory-excitatory synaptic strength (Fig. S1). This result represents a pivotal validation, indicating that gradient-informed heterogeneity allows the model to authentically simulate the physiological constraints where local E-I interactions shape the global aperiodic landscape of brain activity. Ultimately, the TS-DTB transcends mere data fitting to offer a biophysically interpretative platform linking microscale parameters to macroscale signal properties.

While the synergistic benefits of multimodal constraints were evident, it is imperative to ensure that these improvements are robust against the choice of brain atlases. To verify the efficacy of the TS-DTB in reconstructing empirical spatiotemporal dynamics, we assessed its generalization capability using the fine-grained Brainnetome 246 atlas (incorporating subcortical structures) and the high-resolution Schaefer 500 and 1000 cortical parcellations (Fig. S3). Importantly, the mechanistic correlation between the E-I ratio and the aperiodic exponent was consistently preserved regardless of parcellation granularity, confirming that the framework captures robust and invariant biological principles. Furthermore, we evaluated the impact of individualized EEG forward modeling in mapping source-level neural activity to sensor-level signals. While standardized templates serve as a viable approximation in scenarios lacking subject-specific structural MRI, we demonstrated that integrating individualized leadfields significantly enhances reconstruction precision. By introducing personalized anatomical priors, this approach not only minimized spectral relative errors but also more accurately reproduced global nonlinear markers such as metastability and synchrony. Specifically, individualization yielded substantial quantitative improvements in predicting the temporal properties of EEG microstates, including coverage, duration, and transition probabilities (Fig. S4). Collectively, these validations confirm that the TS-DTB extracts reliable biological properties independent of spatial definitions, and that integrating personalized priors further optimizes the reconstruction fidelity of brain dynamics.

### Deciphering E-I imbalance and spectral slowing

Having established the optimal modeling pipeline in the healthy young cohort, we further applied the TS-DTB to an independent AD cohort (Tables S5-S6) to assess its translational capacity (Fig. 4). The models exhibited high fitting accuracy across all groups, including healthy controls, pre-treatment, and post-treatment AD patients, maintaining consistently low REs across a multidimensional suite of temporal statistics, entropy, and spectral features (Fig. 4b, e). Crucially, the TS-DTB models successfully recapitulated macroscopic signatures of cognitive states. It accurately replicated the global explained variance and transition probabilities of empirical EEG microstates (Fig. 4c), and closely matched the empirical characteristics of Lempel-Ziv complexity (LZC). Linear fits for LZC yielded highly significant correlations (*r*=0.837, *R*^*2*^=0.701; *r*=0.789, *R*^*2*^=0.622; and *r*=0.882, *R*^*2*^=0.777; all *p* < 0.001), successfully reproducing the abnormally elevated LZC observed in pre-treatment AD patients compared to controls, as well as the trend of complexity normalization following neuromodulation (Fig. 4d). Spectral analysis revealed that the TS-DTB models accurately captured the pathological slowing characteristic of AD, specifically an increase in low-frequency theta and a decrease in alpha power (Fig. 4f, g)^33,34^. Furthermore, the simulated aperiodic exponents and offsets accurately mirrored the profound empirical downward trends observed in AD patients (e.g., empirical exponent decreased from 1.271 ± 0.015 to 1.207 ± 0.017, *p* < 0.001; simulated exponent decreased from 1.397 ± 0.004 to 1.320 ± 0.004, *p* < 0.001, Fig. 4h)^35^. These precisely captured aperiodic features serve as reliable macroscopic markers for underlying E-I balance shifts associated with cognitive decline and acute post-stimulation normalization.

**Fig. 4.**
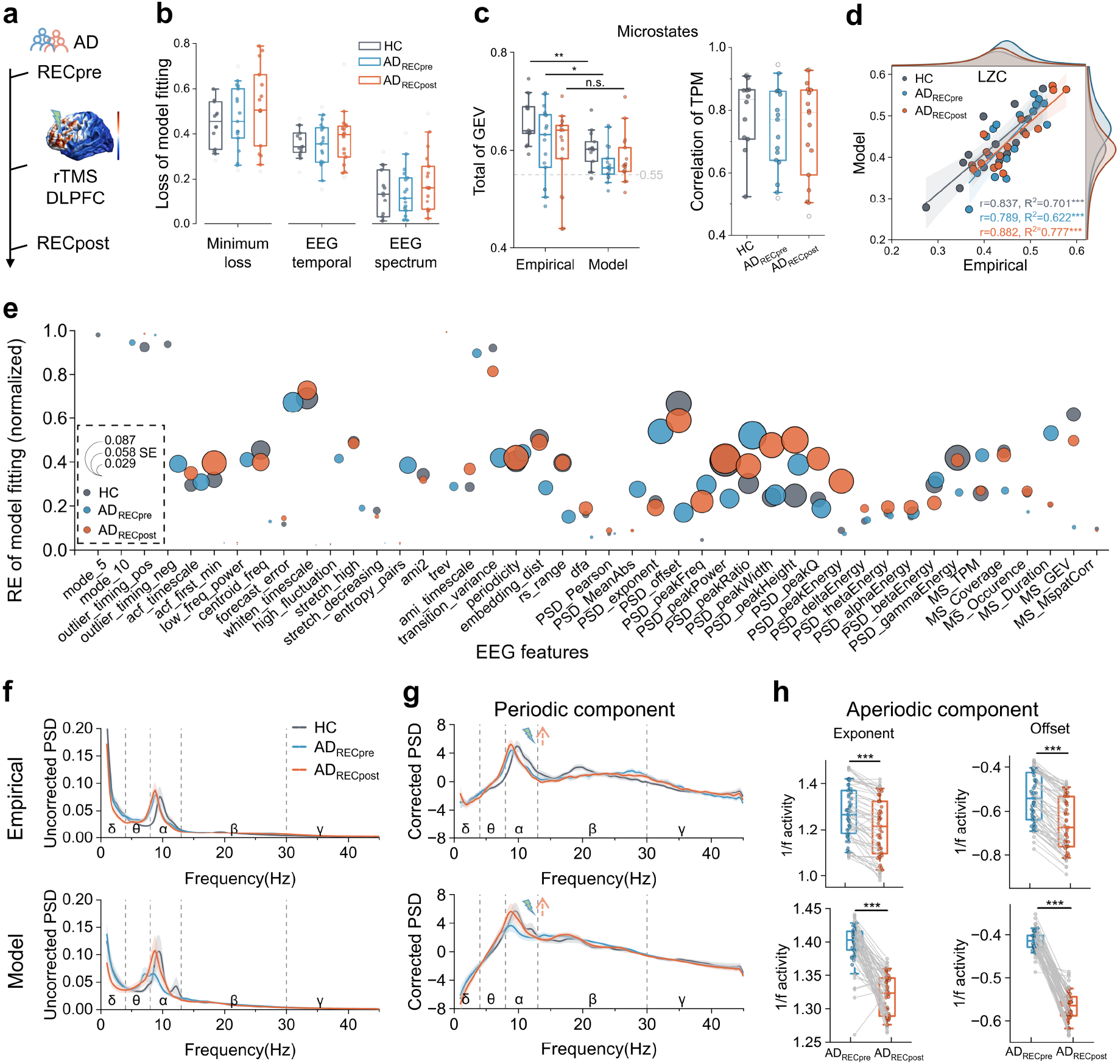
Application of the TS-DTB in studying the cognitive mechanisms of AD. **(a)** Schematic of AD data acquisition undergoing rTMS. **(b)** Global fitting performance. Minimized loss function values demonstrate good and consistent fitting accuracy across HC, pre-treatment AD (AD_RECpre_), and post-treatment AD (AD_RECpost_). **(c)** Microstate dynamics. Total global explained variance (left) and transition probability matrix correlations (right) between empirical and simulated data. **(d)** Normalization of neural complexity. Scatter plots correlating empirical and simulated Lempel-Ziv complexity (LZC), reproducing the elevated complexity in AD and its subsequent normalization post-rTMS. Solid lines indicate linear regression fits and shaded bands represent 95% confidence intervals. **(e)** Feature-wise REs for the evaluated EEG feature set (circle centers indicate the mean RE and diameters are proportional to the SE). **(f)** Spectral reconstruction. Empirical (top) and simulated (bottom) PSD, capturing the pathological slowing in AD and spectral recovery post-rTMS. Solid lines represent the group mean and shaded areas represent the SE across subjects. **(g)** Periodic component analysis. Recovery of alpha peak power post-stimulation. **(h)** Aperiodic component analysis. Across panels (b, c, g, h), boxplots indicate the median (center line) and IQR (box limits), whiskers represent 90% confidence intervals, and dots denote biologically independent individuals (n = 14 HC, n = 17 AD_RECpre_, and n = 17 AD_RECpost_). Significance was assessed via two-sided independent samples t-tests (HC vs. AD_RECpre_) or paired t-tests (AD_RECpre_ vs. AD_RECpost_) (\**p* < 0.05, \*\**p* < 0.01, \*\*\**p* < 0.001).

To dissect the circuit-level mechanisms associated with this cognitive decline, we compared the model parameters and local dynamics between healthy controls and pre-treatment AD patients (AD_RECpre_) (Fig. 5). Crucially, because microstate characteristics were strictly held out from the parameter optimization process, their successful reproduction serves as a robust independent validation of the model’s emergent macroscopic dynamics. The TS-DTB models effectively captured the alterations in microstate temporal characteristics, notably, the deviant duration and coverage of specific microstate classes in the simulated AD brain indicated abnormal frontoparietal activity (Fig. 5a). Functional connectivity analysis demonstrated that the model captured frequency-specific network disruptions, particularly the pathological enhancement of delta-band connectivity and the breakdown of alpha-band connectivity within frontoparietal networks (Fig. 5b, c). At the local circuit level, AD patients exhibited a widespread neural slowing phenomenon, characterized by significantly reduced model-simulated firing rates and inferred E-I ratios across the majority of functional subnetworks (Fig. 5d)^36,37^. Mechanistically, principal component analysis (PCA) of the parameter space revealed a distinct separation between the healthy and AD dynamic states (Fig. 5e). The primary dynamic axis (PC1, explaining 62.7% of the variance) highlighted marked differences primarily influenced by synaptic parameters. To assess parameter stability and identifiability, we validated our framework using 10 independent random initializations. The optimization consistently converged to phenotype-distinct manifolds, achieving an 80.32% phenotype identification accuracy (Fig. S7). Crucially, despite cross-run variations, the core parameter shifts (decreased *c*_*EE*_ and *c*_*IE*_, alongside increased *Ib* and σ in AD) were consistently preserved, aligning with our primary findings. This suggests that the inferred parametric signatures maintain reasonable stability and biological consistency. Furthermore, to validate that the inferred mechanistic parameters (e.g., E-I ratios) are not trivial byproducts of specific optimization targets, we additionally performed a feature hold-out analysis by independently re-optimizing the models using exclusively temporal or exclusively spectral constraints (Fig. S8). However, further mapping of parameter shifts to the E-I ratio indicated that the transition from health to disease was computationally associated with alterations in background input (*Ib*) and stochastic noise levels (σ), rather than structured synaptic coupling alone (*c*_*EE*_ and *c*_*IE*_) (Fig. 5g). Interestingly, this model-inferred sensitivity to noise and background input was most pronounced in low-gradient networks (blue points in Fig. 5g). This finding aligns with the synaptic failure hypothesis, suggesting that as amyloid pathology compromises intrinsic connectivity, the AD brain becomes increasingly susceptible to random fluctuations, resulting in the observed spectral slowing and functional fragmentation.

**Fig. 5.**
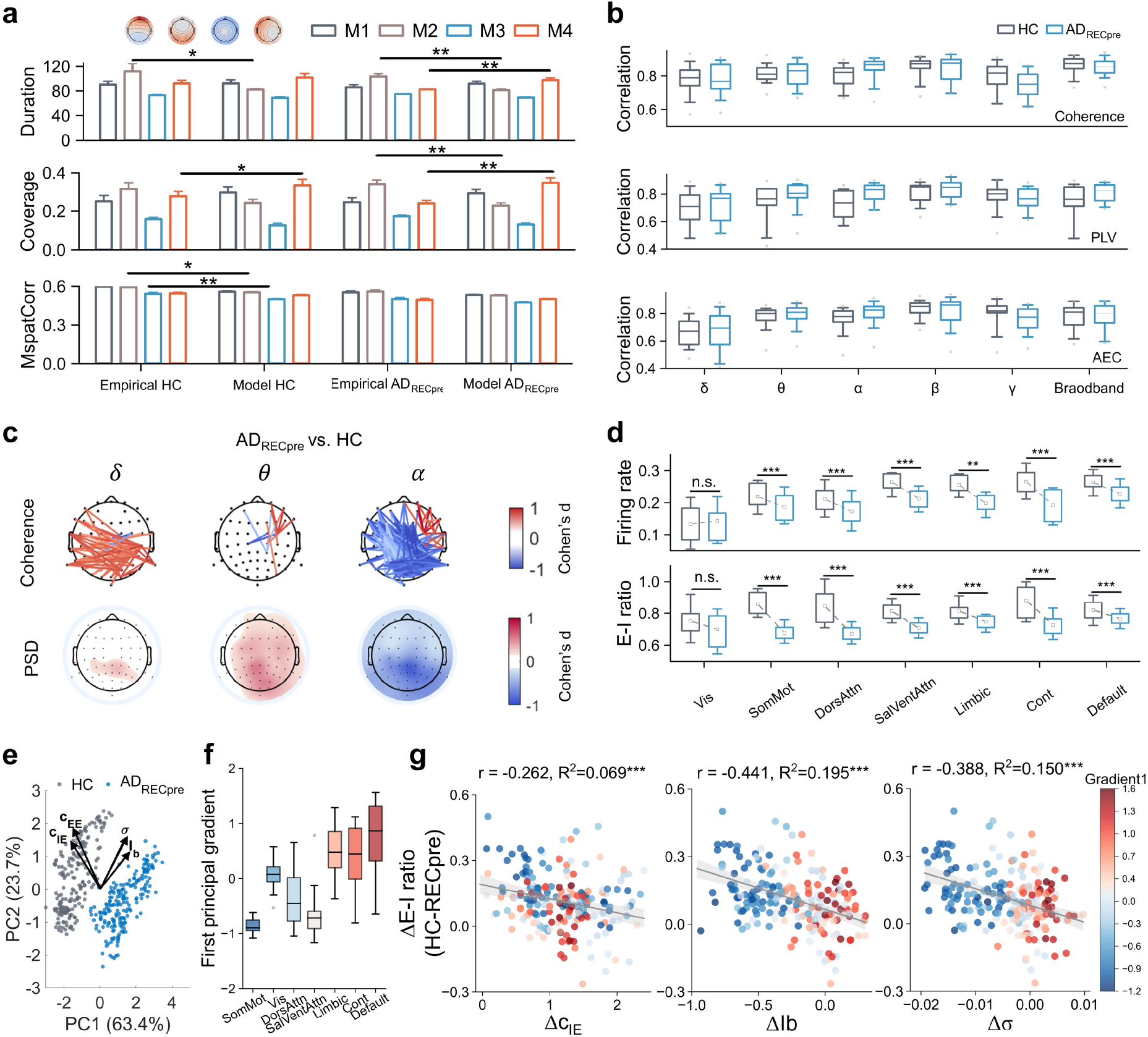
Neurophysiological mechanisms of cognitive decline in AD. **(a)** Microstate temporal characteristics. Comparison of duration, coverage, and spatial correlation of the four canonical microstates between HC and pre-treatment AD (AD_RECpre_). **(b)** EEG FC fitting. Boxplots of correlations between empirical and simulated FC matrices (Coherence, PLV, and AEC) across filter frequency bands. **(c)** Topographic maps of pathological features. Cohen’s d maps illustrating frequency-specific connectivity disruptions and PSD energy alterations in AD. (d) Local circuit dysfunction. Region-wise reductions in simulated firing rates and E-I ratios across functional subnetworks in AD. **(e)** Principal component analysis (PCA). Dynamic axis separation between HC (grey) and AD (blue) derived from the model parameter space (*c*_*EE*_, *c*_*IE*_, *Ib*, σ**). (f)** Functional gradient distribution in HC. Boxplot of the first principal gradient across subnetworks. **(g)** Parametric correlates of cognitive decline (n = 200 cortical regions). Scatter plots correlating parameter shifts (ΔParameter) with E-I ratio alterations. The correlation with changes in background input (Δ*Ib*) and noise (Δσ), particularly in low-gradient regions (blue points) indicates a background-associated pathological regime. For all boxplots (a, b, d, f), the center line indicates the median, box limits represent the IQR, whiskers show 1.5 × IQR, and dots denote individuals in (a, b) (n = 14 HC, n = 17 AD) or cortical regions in (d, f) (n = 200 regions averaged across n = 14 HC and n = 17 AD individuals). Significance was assessed via two-sided independent samples t-tests (\**p* < 0.05, \*\**p* < 0.01, \*\*\**p* < 0.001).

In additional simulations, we observed similar phenomena of E-I imbalance and spectral slowing in a mild cognitive impairment (MCI) cohort (Tables S7-S8 and Figs. S5-S6). Furthermore, to verify cross-site and cross-device generalizability, we applied our modeling pipeline to the independent external BrainLat cohort (Fig. S9). The framework consistently replicated the AD spectral abnormalities and low-dimensional parametric manifold separation between HC and AD. Although scanner site, age, and education introduced systematic baseline variances, disease diagnosis still accounted for independent variance across core parameters. Crucially, the spatial topography of regional parameter associations with cognitive scores for *c*_*EE*_ and *Ib* remained highly consistent across the two cohorts. Consistent with previous experimental findings^31,32,38,39^, these results underscore the cross-dataset robustness of the TS-DTB, demonstrating its generalizable capacity to capture key spatiotemporal signatures of cognitive decline.

### Model-inferred parametric reconfiguration and putative E-I rebalancing underlying rTMS-induced acute electrophysiological normalization

Beyond deciphering the dynamic signatures of AD, the TS-DTB serves as an in-silico testbed to explore the computational correlates of neuromodulation. Therefore, we utilized the framework to investigate the model-inferred parametric shifts associated with acute EEG biomarker normalization following rTMS (Fig. 6). Macroscopically, rTMS induced a systemic normalization of pathological metastability and synchrony in AD patients^40^. This global realignment toward healthy levels was anchored by a significant, band-specific recovery in alpha dynamics, accompanied by broader compensatory trends across other frequency bands (Fig. 6a). This stabilization was underpinned by a recovery in local neural activity, with model-simulated firing rates and inferred E-I ratios increasing overall, particularly in temporoparietal and occipital regions (Fig. 6b, orange vs. blue). The putative rebalancing of the E-I ratio exhibited a spatially heterogeneous pattern, characterized by a disinhibition effect in stimulated frontoparietal areas (increased E-I ratio) and a compensatory dynamic balance in contralateral regions (Fig. 6b, maps).

**Fig. 6.**
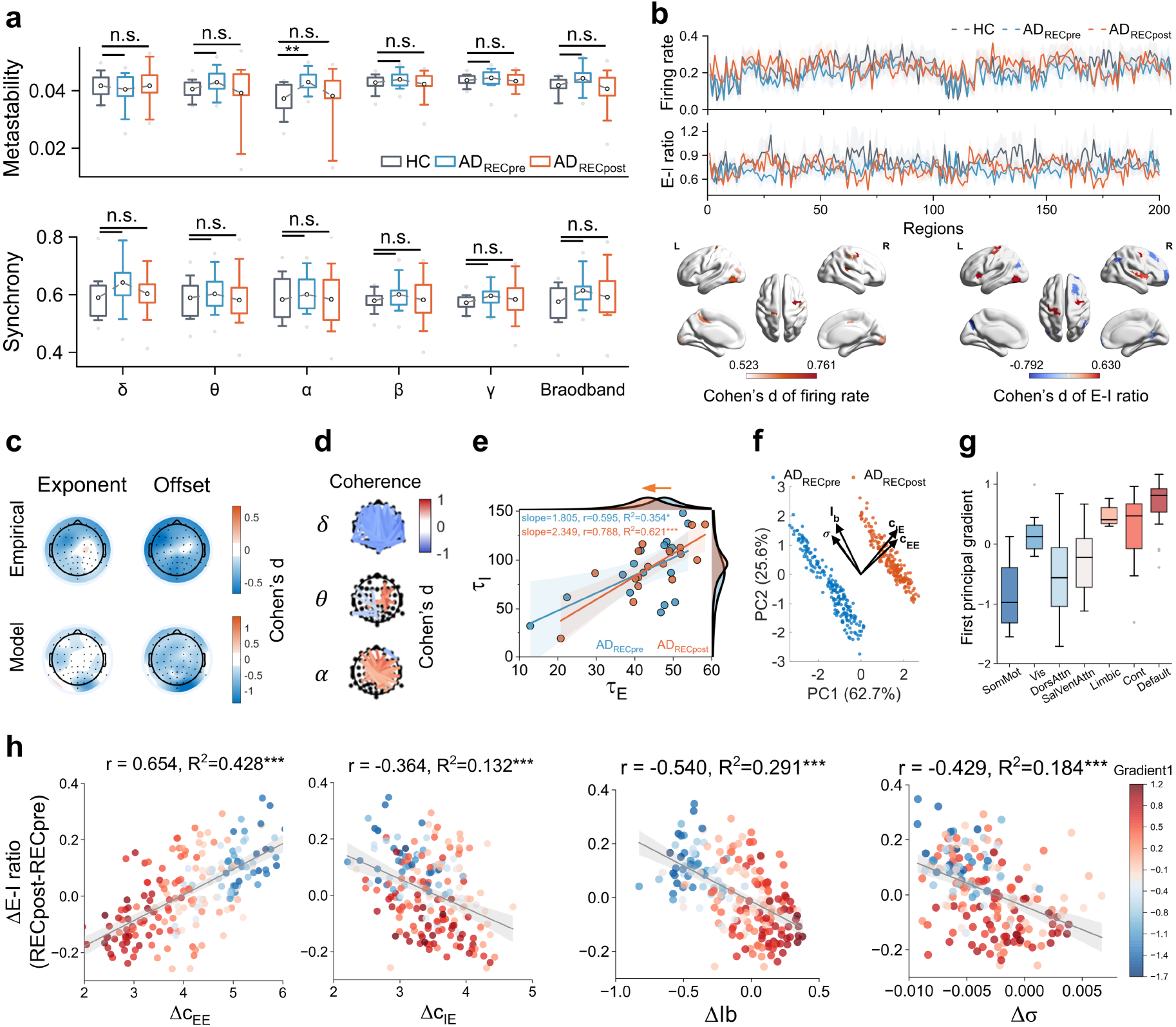
Model-inferred dynamic and parametric reconfiguration induced by rTMS. **(a)** Band-specific normalization of network stability. Normalization of pathological alterations in metastability and synchrony post-rTMS (AD_RECpost_). **(b)** Recovery of local neural activity. Region-wise increases in firing rates and E-I ratios (top), alongside Cohen’s d maps depicting heterogeneous spatial patterns of disinhibition and compensatory balance (bottom). **(c)** Topographic maps. Widespread reductions in aperiodic exponent and offset post-rTMS. **(d)** Network reorganization. Coherence difference maps highlighting the suppression of delta links and re-establishment of frontoparietal theta/alpha connectivity. **(e)** Shift in neural time constants. Significant reduction in the excitatory time constant post-stimulation. **(f)** PCA trajectory of parametric reconfiguration. State shift in parameter space from pre-rTMS to post-rTMS, dominated by synaptic coupling changes. **(g)** Functional gradient distribution in AD patients. **(h)** Gradient-dependent parametric reconfiguration. Scatter plots correlating synaptic parameter shifts (Δ*c*_*EE*_, Δ*c*_*IE*_) with putative E-I rebalancing, showing positive correlations with Δ*c*_*EE*_ in low-gradient regions (blue) and compensatory inhibition (Δ*c*_*IE*_) in high-gradient regions (red) (n=200 cortical regions). For all boxplots and distributions (a, b, e, g), the center line indicates the median, box limits represent the IQR, whiskers show 1.5 × IQR, and dots denote individuals in **(a, e)** (n = 17 AD subjects, pre- vs. post-rTMS) or cortical regions in (b, g) (n = 200 regions averaged across n = 17 AD subjects). Significance was assessed via two-sided paired t-tests (\**p* < 0.05, \*\**p* < 0.01, \*\*\**p* < 0.001).

This normalization was further evidenced by the recovery of functional connectivity, marked by the suppression of pathological delta links and the re-establishment of theta and alpha connectivity in frontoparietal circuits (Fig. 6d)^34^. Additionally, the excitatory time constant τ_*E*_ significantly decreased post-stimulation (*p* < 0.001, Fig. 6e), reflecting faster neural processing. Critically, parameter space analysis revealed that, unlike the background-associated differences between HC and AD_RECpre_, the rTMS-induced electrophysiological normalization was computationally associated with model-inferred synaptic reconfiguration. The PCA trajectories showed a state shift dominated by changes in excitatory and inhibitory synaptic coupling (Fig. 6f). Correlating these parameter shifts with putative E-I rebalancing unveiled a sophisticated, hierarchical pattern of parametric reconfiguration (Fig. 6h)^41,42^. Specifically, in low-gradient regions, the increase in the inferred E-I ratio was strongly positively correlated with increased excitatory coupling (Δ*c*_*EE*_>0). These regions, being relatively spared from AD pathology, exhibited a pronounced parametric shift, potentially acting as a bottom-up compensatory energy source for cortical reactivation. In contrast, high-gradient transmodal regions showed a weaker coupling between Δ*c*_*EE*_ and E-I shifts, frequently accompanied by compensatory adjustments in inhibition (Δ*c*_*IE*_>0). This suggests a homeostatic mechanism where vulnerable core hubs prioritize stability by balancing excitation with inhibition, thereby preventing runaway excitation while integrating the compensatory drive from the periphery (the correlations between parameter shifts and firing rate changes are shown in Fig. S10). Collectively, these model-based interpretations suggest that rTMS induces acute electrophysiological normalization associated with a gradient-dependent parametric reorganization^41,42^, recruiting intact lower-order networks to support the electrophysiological normalization of the global system.

To test whether the model-inferred parametric alterations are computationally sufficient to generate AD-like abnormalities, we performed a parameter perturbation simulation on the individualized HC models along the empirical group-difference vector (Fig. 7a). Macroscopically, progressively shifting the HC models toward the AD-like background abnormality induced pronounced spectral slowing in the periodic PSD component, characterized by a leftward frequency shift and progressive attenuation of the dominant alpha peak alongside elevated low-frequency power (Fig. 7b). Topographic effect-size mapping and band-specific quantification confirmed that negative perturbation scales increased delta power (0.490 to 0.521, *p* < 0.001) and theta power (0.294 to 0.326, *p* < 0.001), while reducing alpha power (0.591 to 0.408, *p* < 0.001) predominantly over posterior cortices (Fig. 7c, d). At the local circuit level, this perturbation trajectory monotonically reduced both the model-inferred E-I ratio (0.828 to 0.722, *p* < 0.001; Fig. 7e) and regional excitatory firing rates (0.230 to 0.197, *p* < 0.001; Fig. 7f), whereas positive scaling reversed these trends. Additional perturbation analyses across the noise parameter are provided in Fig. S11.

**Fig. 7.**
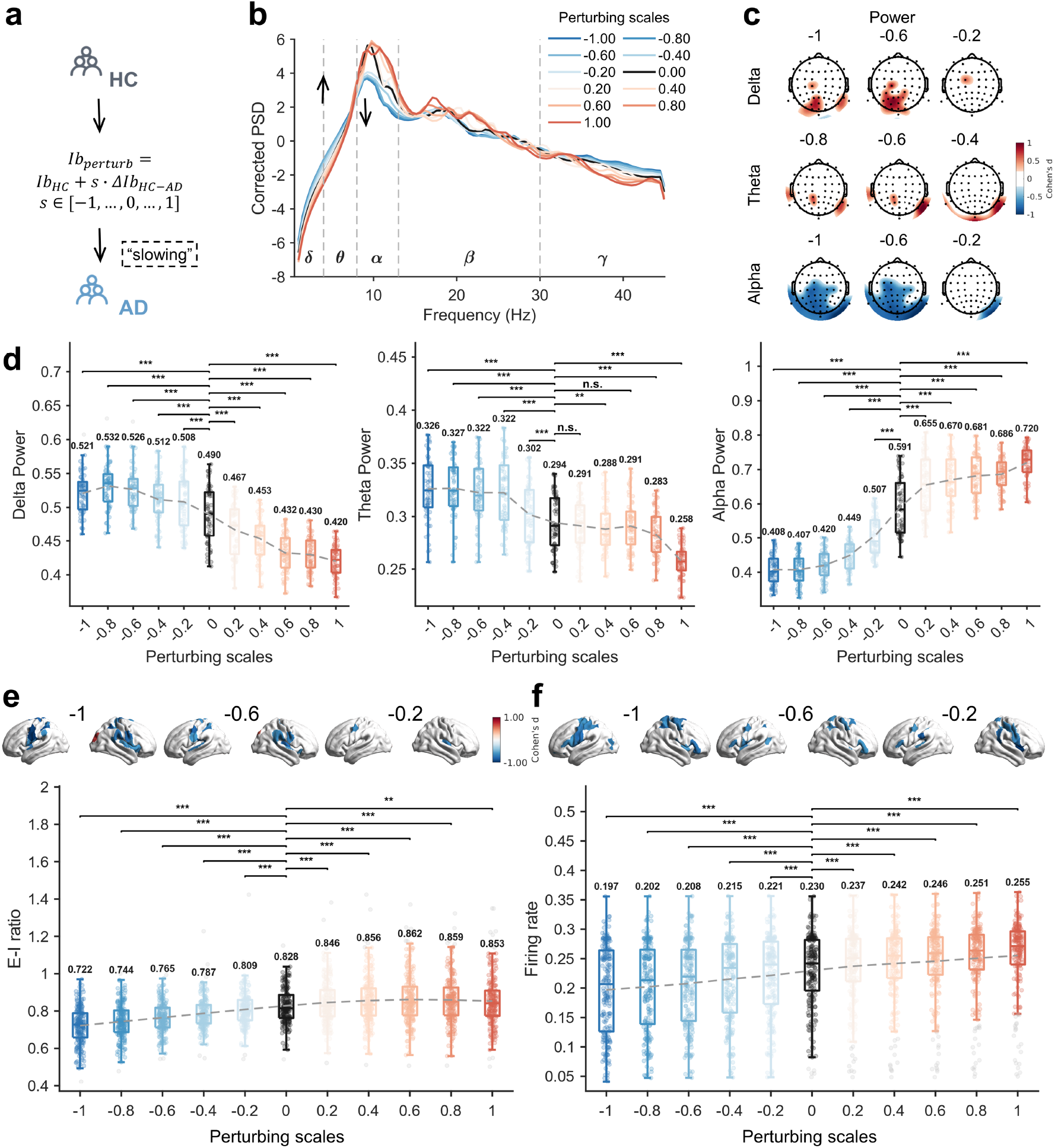
Parameter perturbation in healthy control models induces AD-like spectral slowing and simulated E-I reductions. **(a)** Schematic of the perturbation paradigm. Individualized HC models (n = 14) were systematically perturbed along the empirical group-difference vector (Δ*Ib* _*HC − AD*_) using scaling factors *s* ∈ [−1.0_*k*_1.0] (*Ib*_*perturb*_ = *Ib*_*HC*_ + *s* · Δ*Ib* _*HC − AD*_), where *s* = 0 denotes the unperturbed baseline and s < 0 shifts the model toward the AD regime. **(b)** Channel-averaged periodic power spectral density (Corrected PSD) across perturbation scales, illustrating progressive alpha peak attenuation and leftward shift alongside elevated low-frequency (Delta, Theta) power as s decreases from 0 (black curve) to -1.00 (dark blue curve). **(c)** Topographic Cohen’s d maps comparing perturbed states *s* ∈ [−1.0, −0.2] against baseline (*s* = 0) for delta, theta, and alpha band power. **(d)** Channel-wise power distributions in the delta, theta, and alpha bands across perturbation scales. **(e, f)** Region-wise distributions of the model-inferred E-I ratio (e) and simulated excitatory firing rates (f) across perturbation scales, accompanied by cortical surface Cohen’s d maps for negative scales (*s* = -1.0, -0.6, -0.2 vs. baseline *s* = 0). For all boxplots in (d-f), the central line indicates the median, box limits represent the interquartile range (IQR), whiskers extend to 1.5 × IQR, numerical labels indicate group means, and dots denote EEG channels in (d) (n = 61 channels) or cortical regions in (e, f) (n = 200 regions) averaged across n = 14 biologically independent HC subjects. Statistical significance relative to the baseline (*s* = 0) was assessed using two-sided paired t-tests (\**p* < 0.05, \*\**p* < 0.01, \*\*\**p* < 0.001; n.s., not significant).

To evaluate the forward-predictive capacity of the TS-DTB framework, we performed a physics-constrained simulation of rTMS intervention on the individualized baseline AD models (AD_RECpre_) (Fig. 8a). We utilized SimNIBS to derive the realistic cortical distribution of the induced electric field (eField) targeting the left DLPFC (F3) and mapped it as a localized spatial perturbation onto the background input (*Ib*) and noise (σ) parameters, governed by global scaling factors and an exponential temporal decay constant (Fig. 8a). By tuning these low-dimensional intervention hyperparameters via Bayesian optimization, the baseline models forward-generated simulated post-intervention EEG dynamics. Using the post-rTMS model state (AD_RECpost_) as a reference, the virtual intervention yielded emergent macroscopic EEG alterations that consistently aligned with the post-rTMS trajectory across both uncorrected and aperiodic-corrected power spectra (Fig. 8b, c). Specifically, the eField model (eField-AD) captured the suppression of low-frequency delta power, the enhancement of alpha power, and the reduction of aperiodic exponent and offset, alongside computationally inferred increases in local E-I ratios and firing rates (Fig. 8d). To test spatial specificity, we constructed a spatial-autocorrelation-preserving Moran null model by spatially scrambling the eField topography. Notably, while the null model partially mimicked the directional trends in narrowband delta and alpha power as well as E-I ratio due to global perturbation energy, it failed to increase the firing rate and induced opposite changes in the aperiodic exponent and offset (Fig. 8d). These simulated results reflect that the acute normalization of both oscillatory and aperiodic electrophysiological signatures is associated with the realistic spatial topography of the rTMS-induced electric field, highlighting the utility of the TS-DTB framework as an in-silico testbed for neuromodulation.

**Fig. 8.**
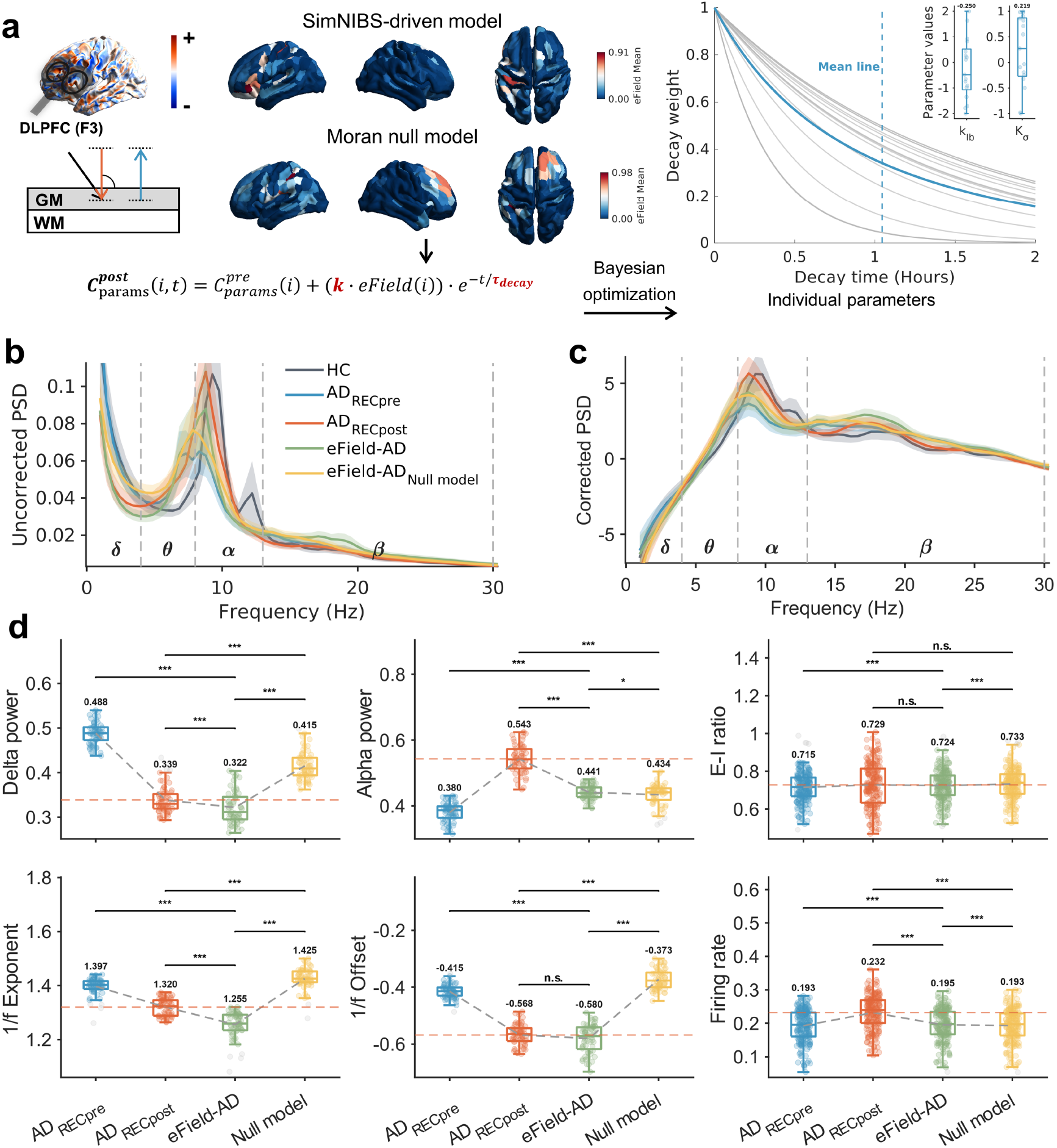
Forward simulation of rTMS intervention via SimNIBS-driven digital twins. **(a)** Workflow of the virtual intervention simulation. The realistic electric field (eField) targeting the left DLPFC (F3) generated by SimNIBS (top maps) and a spatial-autocorrelation-preserving Moran null model (bottom maps) were mapped onto cortical regions to modulate baseline parameters via scaling factors (k) and an exponential temporal decay function 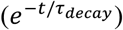 optimized through Bayesian optimization. Right: individual (grey) and group-mean (blue; vertical dashed line indicates mean decay time) decay weight trajectories over 2 hours, with inset boxplots of optimized scaling parameters (n = 17 AD subjects). **(b, c)** Comparison of uncorrected (b) and aperiodic-corrected (c) PSD profiles across HC, baseline AD (AD_RECpre_), post-rTMS target AD models (AD_RECpost_), the realistic eField-driven model (eField-AD), and the null model (eField-AD_Null model_). Solid curves represent the group mean and shaded areas represent the SE. **(d)** Quantitative evaluation across delta power, alpha power, E-I ratio, 1_2_*f* exponent, offset, and firing rate (horizontal dashed red lines denote mean AD_RECpost_ reference level). Unlike the Null model, which fails to elevate firing rates and induces opposite trajectories in aperiodic components, eField-AD consistently shifts all oscillatory, aperiodic, and model-inferred metrics toward the AD_RECpost_ target. For all boxplots in (d), the center line indicates the median, box limits represent the IQR, whiskers extend to 1.5 × IQR, and dots denote channel-wise values (n = 61 EEG channels averaged across n = 17 biologically independent AD subjects for spectral and aperiodic metrics) or region-wise values (n = 200 cortical regions averaged across n = 17 biologically independent AD subjects for E-I ratio and firing rate). Significance was assessed via two-sided paired t-tests (\**p* < 0.05, \*\**p* < 0.01, \*\*\**p* < 0.001; n.s. indicates not significant).

## Discussion

In this study, we developed the TS-DTB modeling framework to address the fundamental challenge of reconciling disparate spatiotemporal scales in individualized brain dynamics modeling. By implementing a source-to-sensor optimization strategy, the TS-DTB effectively integrates individualized hemodynamic and electrophysiological constraints, resolving the parameter coordination inherent in single-modality approaches^19,23^. Our validation across both healthy and AD cohorts demonstrated that the generated TS-DTB models surpass conventional optimization schemes by simultaneously reconstructing large-scale network topologies and reproducing fast spatiotemporal dynamics^6,22,23^. Specifically, evaluating the framework in a healthy cohort established its general modeling superiority. More importantly, applying it to the AD cohort allowed us to computationally characterize the model-inferred circuit parameters associated with macroscopic cognitive decline and to in-silico track acute electrophysiological normalization.

The capability to dissect these nuanced spatiotemporal mechanisms relies fundamentally on the holistic integration of multimodal data feature constraints, combining fMRI-derived topology, gradient-informed heterogeneity, and individualized EEG forward modeling. Without these constraints, traditional homogeneous models fail to capture the hierarchical complexity of brain plasticity, erroneously collapsing region-specific dynamic responses into a uniform profile^11,27,43,44^. By constraining local circuit heterogeneity, our approach effectively resolves parameter coordination, enabling the precise estimation of region-specific synaptic strengths and background-related parameters. This biophysically grounded fusion leverages the high spatial resolution of fMRI to disentangle the underlying source-level interactions that drive observable sensor-space oscillations. Furthermore, the incorporation of individualized FEM-based leadfields proved indispensable for resolving the spatial ambiguity of EEG sources, ensuring that modeled cortical dynamics accurately project to real-world sensor observations^28^. This multimodal information jointly constrained strategy differs from previous approaches, offering an effective pipeline for reconciling hemodynamic topology with electrophysiological kinetics.

Applying this unified framework to the AD cohort, our model suggests that the macroscopic spectral slowing and abnormal complexity characteristic of the disease are computationally associated with a model-inferred disruption of the local E-I balance^20,40^. Specifically, parameter space analysis in pre-treatment AD patients revealed a gradient-dependent E-I shift computationally associated with aberrant background suppression, which most severely afflicts low-gradient regions. This aligns robustly with the synaptic failure hypothesis, suggesting that as amyloid pathology compromises intrinsic connectivity, neural circuits become increasingly susceptible to random fluctuations, leading to functional network fragmentation and the dissolution of metastable microstates^32,38^. Similarly, consistent E-I imbalance patterns were also observed in an independent MCI cohort, reinforcing the generalizability of our framework^45^. By directly mapping and linking macroscopic EEG abnormalities to specific microcircuit parameters, the TS-DTB models establish the model-derived E-I ratio as a computational proxy that correlates with cognitive states. This capacity to inversely map sensor-level phenotypes to source-level parametric alterations provides a critical theoretical platform for dissecting the neural basis of cognitive decline and post-stimulation normalization and the design of future neuromodulation strategies.

Another contribution of this work lies in leveraging the TS-DTB as an in-silico testbed to computationally characterize rTMS-induced acute electrophysiological normalization. Challenging the conventional assumption that neuromodulatory effects are strictly confined to the stimulation target, our analysis suggested a sophisticated, gradient-dependent pattern of putative E-I rebalancing characterized by a global synergy of peripheral drive and core homeostasis^41,42^. Specifically, we observed that simulated acute EEG normalization is computationally associated with extensive model-inferred synaptic reconfiguration within low-gradient sensorimotor networks. This actively recruits structurally intact lower-order regions as an energy reservoir to drive global cortical re-awakening, effectively compensating for the inertia of damaged higher-order hubs (e.g., the default mode network)^11,26^. Concurrently, to maintain homeostatic stability within the pathological transmodal core, high-gradient regions exhibited a distinct mechanistic profile characterized by profound background suppression alongside a compensatory increase in inhibition. Ultimately, this hierarchical reorganization provides a biophysically grounded theoretical framework for understanding how focal stimulation induces global electrophysiological normalization, highlighting that effective neuromodulation relies on the coordinated rebalancing of E-I ratios across the entire cortical hierarchy^46,47^. While the post-rTMS spectral and aperiodic improvements represent robust empirical observations, the accompanying gradient-dependent shifts in synaptic coupling and E-I ratios represent macroscopic model-based interpretations rather than direct measurements of cellular synaptic plasticity. Establishing definitive causal pathways linking rTMS to microcircuit plasticity and long-term clinical improvement will require future studies integrating longitudinal behavioral assessments with direct neurophysiological recordings.

Several limitations of the current study should be noted. First, the clinical cohorts utilized for our computational modeling are relatively small and not strictly age-matched. This demographic mismatch, arising from recruitment constraints, may introduce age-related confounding effects into the inferred model parameters. Moreover, cross-site heterogeneities in MRI scanners, EEG systems, and demographics inevitably introduce baseline variances into the inferred parameter space, as quantified by our variance partitioning analysis (Fig. S9d). While direct comparisons of absolute parameter magnitudes across sites should therefore be interpreted with caution, our external validation on the BrainLat cohort confirmed that both the low-dimensional manifolds and the cortical topography linking inferred parameters to cognitive scores remain reproducible across heterogeneous environments (Fig. S9a, e).

Looking forward, the TS-DTB presents several promising avenues to further enhance biophysical plausibility and computational efficiency^13^. First, future studies could extend the complexity of the underlying physics by incorporating transmission delays, directed effective connectivity and feedforward inhibition^25,48^. In addition, the biological grounding of parameter heterogeneity could be enriched by integrating a broader spectrum of multiscale priors. Beyond functional gradients, incorporating intrinsic timescales, myelin maps, or transcriptomic profiles could provide multidimensional constraints, yielding a more integrative view of cortical organization^10,49,50^. Finally, while the current study modeled rTMS effects as parameter shifts between pre- and post-states, future work should aim to simulate dynamic plasticity rules directly within the model. Such advancements would enable fully predictive in-silico trials to optimize patient-specific protocols before intervention^51,52^.

In summary, the TS-DTB establishes a generalizable computational platform that effectively bridges the spatiotemporal gap inherent in conventional brain modeling. By synergizing spatial heterogeneity with individualized EEG forward mapping, this approach successfully reconciles the macroscopic topology of hemodynamics with the fast electrophysiological signatures of local microcircuits. Crucially, the capacity of TS-DTB to inversely map non-invasive sensor-level features to underlying mechanistic parameters offers a transformative lens for dissecting neuropathology. While demonstrated here in the context of AD and rTMS, this biologically grounded methodology provides a universal foundation for decoding multiscale circuit disruptions across a broad spectrum of neuropsychiatric disorders. Ultimately, the TS-DTB transcends data-driven descriptive neuroimaging, paving the way for in-silico clinical trials and the realization of model-guided digital research^13,14^.

## Methods

### Multimodal neuroimaging datasets

This study utilized four independent cohorts, with all protocols approved by institutional ethics committees and informed consent obtained. All experimental procedures involving human participants were conducted in accordance with the Declaration of Helsinki. Written informed consent was obtained from all participants prior to enrollment. The healthy cohort consisted of 25 young adults (13 females, age 24.2 ± 1.2 years) recruited via campus advertisements (with standard compensation) at the University of Electronic Science and Technology of China, with ethical approval granted by the Ethics Committee of the University of Electronic Science and Technology of China. The Alzheimer’s disease (AD) clinical cohort, recruited from outpatient clinics at Shenzhen People’s Hospital, included 17 patients (14 females, age 65.5 ± 9.5 years) and 14 healthy controls (HCs, 12 females, age 56.4 ± 5.3 years; Tables S5–S6), with ethical approval granted by the Medical Ethics Committee of Shenzhen People’s Hospital. The mild cognitive impairment (MCI) clinical cohort, recruited from the Fourth People’s Hospital of Chengdu, comprised 15 patients (8 females, age 64.7 ± 5.2 years) and 16 age-matched HCs (12 females, age 62.6 ± 5.5 years; Tables S7–S8), with ethical approval granted by the Ethics Committee of the Fourth People’s Hospital of Chengdu. An independent external public dataset (BrainLat) was also incorporated, comprising 15 AD patients (10 females, age 77.60 ± 9.19 years) and 10 HCs (5 females, age 74.40 ± 3.03 years; Tables S9–S10), which was acquired under prior institutional ethical approvals as part of the open-access BrainLat^53^. The detailed information on data acquisition and preprocessing is available in the Supplementary Information.

### Large-scale brain dynamics model

#### Neural mass model

We simulated macroscopic brain activity using a network of coupled Wilson-Cowan oscillators, which provides a biologically grounded, nonlinear description of regional population dynamics^54^.

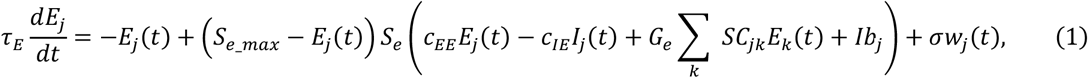

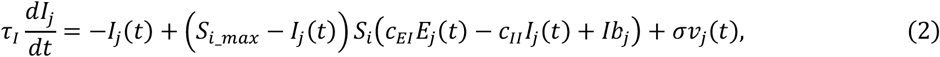

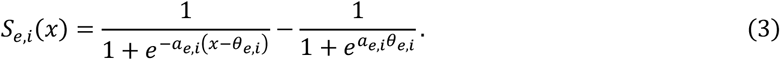

Specifically, the state of the *j*-th brain region at time *t* is defined by the normalized firing rates of its excitatory (*E*_*j*_(*t*)) and inhibitory (*I*_*j*_(*t*)) populations. The temporal evolution of these populations is governed by their respective time constants τ_*E*_ and τ_*I*_. The input-output relationship *S*_*e,i*_(*x*) for each population is modeled via a sigmoidal transfer function, characterized by a maximum activity 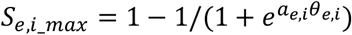, a mean firing threshold θ_*e,i*_ and variation slope *a*_*e,i*_. The dynamics of the excitatory pool ensemble are driven by local synaptic connection strength interactions between the excitatory and inhibitory populations, long-range inputs coupling the structural connectivity matrix *SC*_*jk*_ (which is substituted by the optimized effective connectivity *EC*_*jk*_ in the fMRI-constrained optimization models) from different populations with global coupling strength *G*_*e*_, and the background input *Ib*_*j*_. Additive noise represents the stochastic perturbation input to the system with the functions *w*_*j*_(*t*) and *v*_*j*_(*t*), which are derived from a normal Gaussian distribution with a standard deviation σ. Other constants in the model were biologically derived as described in previous work^25,54^. Numerical simulations were performed using the Euler method with an integration step of 0.1 ms, initializing values *E*_*j*_(0) = *I*_*j*_(0) = 0.1. Following the exclusion of a 20 s transient period to ensure dynamic stabilization, the *E*(*t*) were downsampled for subsequent multimodal forward projections.

#### Multimodal forward modeling

Simulated BOLD signals were derived from the excitatory firing rate signals using the standard Balloon-Windkessel hemodynamic model, which mathematically couples neural firing rate (*E*) to changes in blood flow, volume, and deoxyhemoglobin content^55^. The hemodynamic changes at node *j* were as follows:

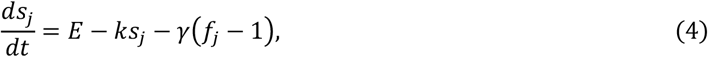

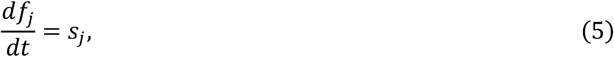

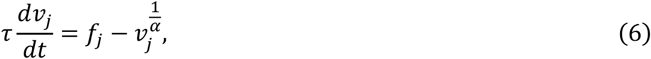

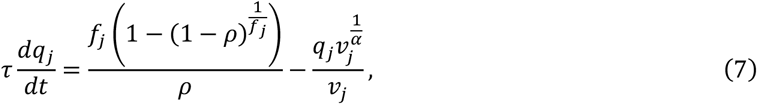

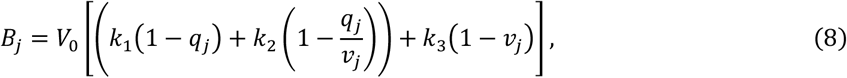

where *B*_*j*_ represents the simulated BOLD signal from node *j*. The resting blood volume fraction *V*_0_ = 0.02, *k*_1_ = 7ρ, *k*_2_ = 2, and *k*_3_ = 2ρ − 0.2, other parameters were taken from^29^.

Concurrently, to reconstruct sensor-level electrophysiology, the *E*(*t*), representing source-space current dipoles, were projected to the scalp utilizing the leadfield (*LF*) computed via the FEM-based head models^56^. This precise spatial mapping explicitly accounts for realistic volume conduction effects across distinct tissue compartments, yielding the simulated scalp EEG signals (*S*_*EEGS*_):

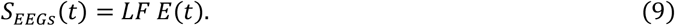

### Model optimization

#### Inter-regional coupling calibration in the first stage

We aimed to refine the source-space effective coupling (EC) matrix to accurately reflect macroscopic hemodynamic interactions. Adopting established structural-functional iterative schemes^8^, we employed an fMRI-based optimization procedure (fMRIOpt) that quantified the discrepancy between the empirical and simulated FC. The loss function minimized both the Pearson correlation and the Kolmogorov-Smirnov distance (KSD) between the respective FC distributions:

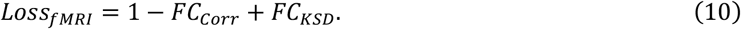

To achieve convergence, the EC matrix was iteratively updated by adjusting an adaptive learning rate *Lr*_*x*_ with decay^8^. It should be noted that our optimization procedure allows for negative entries in the updated EC matrix. Because our model architecture restricts long-range inputs to target the local excitatory populations directly, these negative effective weights serve as a net-input strategy to model inter-regional suppression. This functional abstraction allows the model to efficiently capture both cooperative and competitive dynamics across large-scale brain networks^57^. The globally optimized EC matrix was subsequently fixed for the second stage. Specifically, this optimized EC replaced the empirical SC in all fMRI-constrained models (fMRIOpt and TS-DTB), whereas models without fMRI optimization (noOpt and EEGOpt) retained the original SC:

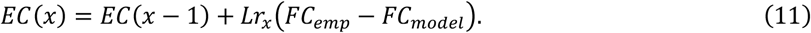

In this stage, we fix the global coupling strength *G*_*e*_ = 1. The direction of change in FC correlation was quantified as Δ*Corr* = *FC*_*Corr*_(*x* + 1) − *FC*_*Corr*_(*x*). The optimization was performed using iterations *X* = 30, with an initial learning rate *Lr*_0_ = 0.025, decay rate *Dr* = 0.02, increase factor *If* = 0.25, decrease factor *Df* = −0.1, and the *Lr*_*x*_ in the *x*^th^ iteration adjustments were restricted to the range *Lr*_*x*_ ∈ [0.001,0.05]. The adaptive learning rate evolved according to

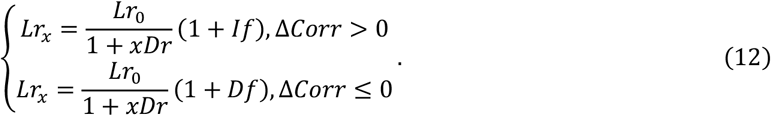

### Intra-regional dynamics optimization in the second stage

Building upon the calibrated network backbone, the second stage optimized region-specific kinetic parameters to reproduce fast, sensor-space EEG dynamics^11,27^. Specifically, the macroscale organizational structure of the brain was characterized using the first two functional connectivity gradients (denoted as *Grad*_1_ and *Grad*_2_) derived via the BrainSpace toolbox^58,59^. These gradients provided fMRI-based priors that capture fundamental hierarchical patterns of regional integration and segregation. We modeled regional heterogeneity by parameterizing the inhibitory to excitatory synaptic strength (*c*_*IE,j*_), background input (*Ib*_*j*_), and noise amplitude (σ_*j*_) as linear combinations of these two empirical macroscale gradients. Consequently, each variable was governed by a distinct gradient-weighting profile, defined by linear coefficients (*a*_*k*,_ *b*_*k*,_ *c*_*k*_):

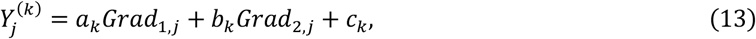

where 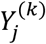 denotes the value of parameter *k* (i.e., *k* ∈ {*c*_*IE*,_ *Ib*, σ} for region *j*. We constructed two ablation models to verify the necessity of gradient-informed priors. The He_Moran_ models replaced empirical gradients with Moran-randomized surrogates, nullifying authentic spatial topography while retaining spatial autocorrelation. Conversely, the Ho models assumed spatial homogeneity, enforcing uniform local kinetic parameters across all regions. During optimization, we sampled the exponent *x* from a uniform prior distribution to define σ = 10_*x*_, ensuring that the noise standard deviation remained strictly positive. The parameter space consisted of gradient-weighting parameters and three globally co-optimized variables, including excitatory time constant (τ_*E*_), inhibitory scaling factor (*r*_*τI*_ here τ_*I*_ = *r*_*τI*_ τ_*E*_), and global coupling (*Ge*). For AD cohort simulations, an additional parameter for excitatory– excitatory synaptic coupling (*c*_*EE*_) was introduced to model rTMS-induced effects (detailed in Table S1).

The EEG-based loss function integrated both temporal and spectral features, which were extracted using the catch22, FOOOF, and standard signal processing pipelines (Tables S2-S3)^60,61^. The discrepancy between simulated and empirical EEG signals was quantified using the median relative errors (REs, normalized to the range of [0,1]) across these feature sets. The temporal and spectral components were equally weighted to formulate the final loss function:

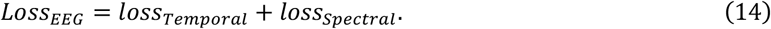

To minimize the loss function, we performed Bayesian optimization using the expected-improvement-plus (EI_+_) acquisition function within a 32-core parallelized environment. The process executed 96 total evaluations (initial seed to Gaussian process guided ratio of 1:3), utilizing an exploration ratio of 0.75.

### Parameter perturbation and SimNIBS-driven rTMS simulation

To evaluate whether the model-inferred parameter shifts are sufficient to induce AD-like abnormalities, we conducted parameter perturbation simulations on the individualized HC models (n = 14) from the AD clinical cohort. First, the empirical group-level difference vector between the HC and pre-treatment AD cohorts was computed across all cortical regions 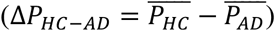. Next, for each baseline HC model, the regional background input parameter was systematically perturbed along this difference vector across 11 equally spaced scaling factors (*P*_*perturb*_ = *P*_*H*C_ + *s* · Δ*P*_*H*C − *AD*_,*s* ∈ [−1.0, 1.0], where *s* = 0 corresponds to the unperturbed HC baseline and negative *s* shifts the parameter profile to the AD pathological regime). All other calibrated parameters were held at their subject-specific baseline values, and simulations were executed at each scaling step to quantify the resulting regional firing rates, model-inferred E-I ratios, and sensor-level EEG spectral properties.

To validate the forward-predictive capacity, we conducted the rTMS intervention simulation on the individualized pre-rTMS AD baseline models (n = 17). First, we utilized SimNIBS to derive the realistic, subject-specific spatial distribution of the electric field (eField) induced by rTMS over the left dorsolateral prefrontal cortex. We then mapped this physical field onto the cortical surface as a spatial modulator of local background input (*Ib*) and noise (σ) parameters. To reflect decay responses, we incorporated a temporal decay function 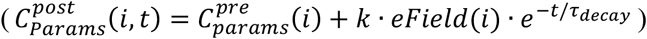, where *k* represents the eField scaling factor and τ_*decay*_ is the decay time constant. By optimizing these low-dimensional global parameters via Bayesian optimization, the models forward-simulated post-rTMS macroscopic EEG dynamics. Furthermore, to test the spatial specificity of the eField, we generated a spatial null model using Moran spectral randomization, which scrambled the realistic eField topography while preserving its spatial autocorrelation.

## Data analysis

### Excitation-inhibition (E-I) ratio

At the microcircuit level, the E-I balance was quantified from the simulated neural activity. The E-I ratio was defined as the ratio between the temporal average of the excitatory and inhibitory firing rates^27^

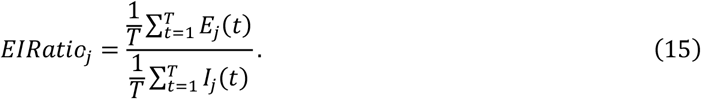

### Network synchrony and metastability

Global dynamical complexity was characterized using two conventional metrics for coupled-oscillator systems: network synchrony and metastability^62^. These quantities derive from the Kuramoto order parameter *R*(*t*), defined as

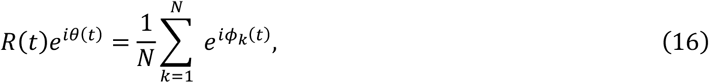

where *ϕ*_*k*_(*t*) is the instantaneous phase of the *k*-th regional signal. Phases were obtained from the analytic signal

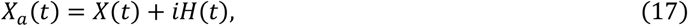

where *X*_*a*_ is the analytic signal, *X* is the original signal, and *H* is the Hilbert transform of *X*. The analytic signal was expressed in polar form as

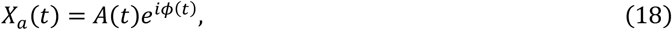

where *A*(*t*) is the amplitude envelope, and *ϕ*(*t*) is the instantaneous phase. Network synchrony was quantified as the temporal mean of the *R*(*t*), reflecting the overall degree of phase alignment across the network over time. Metastability was defined as the standard deviation of the *R*(*t*), indexing the temporal variability of this synchronization, thereby capturing the tendency of the system to continuously explore diverse coordination states.

### Statistical analysis

All correlations were evaluated using the Pearson correlation coefficient (*r*). Differences between empirical and simulated feature distributions, or between distinct clinical groups/conditions, were assessed using paired or two-sample tests as appropriate. Normality was evaluated using the Shapiro-Wilk test. Normally distributed data were analyzed using paired or independent t-tests, whereas non-parametric equivalents (Wilcoxon signed-rank test or Mann-Whitney U test) were applied otherwise. The results were considered significant at \**p* < 0.05, \*\**p* < 0.01 and \*\*\**p* < 0.001. Effect sizes were reported by Cohen’s d.

## Data availability

The external BrainLat dataset^53^ analyzed in this study is publicly available via Synapse (https://www.synapse.org/Synapse:syn51549340/wiki/624187)^63^. The in-house datasets from the healthy young, MCI, and AD cohorts contain sensitive clinical and anatomical information; due to patient privacy regulations and restrictions imposed by the approving institutional ethics committees, these raw datasets are under controlled access and are not publicly deposited. De-identified data are available from the corresponding author (Prof. Daqing Guo) upon request for non-commercial academic research, subject to institutional ethical approval and a formal data use agreement prohibiting participant re-identification. The source data underlying Figs. 1–8 are provided in Supplementary Data 1-8, respectively.

## Code availability

The custom code and the TS-DTB modeling framework used to generate and analyze the results in this study are publicly available on Zenodo (https://doi.org/10.5281/zenodo.23686789)^64^.

## Author contributions

Y.X., D.Y., and D.G. conceived the study and developed the methodology. Y.X., Y.Xu, and Y.C. developed the software and preprocessed the data. R.Z., Y.L., F.W., and X.Z. performed the statistical analysis and curation. D.G., Y.G. and D.Y. provided resources. Y.X. and D.G. wrote the original draft. D.G. and D.Y. acquired funding, supervised the project, and revised the manuscript. All authors reviewed and approved the final version.

