## Supplementary material for "An EEG-fMRI Jointly Constrained Digital Twin Brain and Its Application in Alzheimer’s Disease": SI

### Supplementary Information

#### Data preprocessing and analysis

##### MRI acquisition

Structural (T1w), diffusion-weighted imaging (DWI), and resting-state functional MRI (rs-fMRI) data were acquired using 3T scanners (GE Discovery MR750 for the healthy cohort; Siemens Skyra for both clinical cohorts). High-resolution T1w images were acquired using standard 3D MPRAGE or IR-GRE sequences (voxel size=$1\times1\times1$ $mm^3$). DWI data were collected using single-shot EPI sequences with 64 diffusion directions (voxel size =$2\times2\times4$ $mm^3$). The rs-fMRI data were acquired using gradient-echo EPI sequences with a temporal resolution (TR) of approximately 2000 ms, capturing over 255 volumes for the healthy cohort and 238 volumes for the clinical cohorts per participant.

##### EEG acquisition

Eyes-closed resting-state EEG signals were recorded using 64-channel caps conforming to the international 10-20 system. Recordings utilized ASA-Lab amplifiers (ANT Neuro, Netherlands) for the healthy cohort and LiveAmp 64 systems (Brain Products, Germany) for the clinical cohorts. The sampling rates were 1000 Hz or 5000 Hz, which were subsequently downsampled during preprocessing. Uniquely, the AD group underwent a 30-min session of repetitive transcranial magnetic stimulation (rTMS) targeting the left dorsolateral prefrontal cortex (DLPFC), and then the EEG data were acquired immediately before (RECpre) and after (RECpost) the intervention.

##### Data Preprocessing

We utilized four distinct brain atlases at 1 mm voxel resolution, including the Schaefer 200, 500 and 1000 cortical parcellations[1], and the Brainnetome 246 atlas[2], to define regions of interest (ROIs) for multimodal network and leadfields construction.

##### Structural connectivity (SC)

SC matrices were derived from DWI data using the MRtrix3 toolbox[3]. Following standard denoising and motion/bias field correction, images were registered to the MNI152 template using ANTs. Tissue segmentation was performed using FSL 5ttgen for anatomically constrained tractography (ACT). Fiber orientation distributions were estimated via constrained spherical deconvolution. Whole-brain probabilistic tractography (iFOD2 with ACT) generated 200,000 streamlines per participant. SC matrices were constructed by quantifying the normalized streamline counts connecting each pair of ROIs.

##### fMRI BOLD signals

The rs-fMRI data preprocessing was conducted using DPABI[4]. The fMRI images were slice-time corrected, spatially realigned, normalized to MNI space ($3\times3\times3$ $mm^3$), and smoothed with a 6 mm full-width half-maximum (FWHM) Gaussian kernel. The nuisance signals, including 24 motion parameters and mean white matter and cerebrospinal fluid (CSF) signals, were regressed out. The resulting BOLD time series were bandpass filtered (0.01–0.1 Hz) and z-score normalized for each ROI.

##### Functional connectivity (FC)

For fMRI, FC matrices were constructed by computing Pearson correlation coefficients between all pairs of regional BOLD time series, followed by Fisher's z-transformation to improve normality. For EEG, sensor-level FC was quantified across filter frequency bands (delta to gamma) using three complementary metrics to capture diverse interaction mechanisms: (1) Amplitude envelope correlation

(AEC) measures the correlation of orthogonalized amplitude envelopes; (2) Magnitude-squared coherence quantifies the linear dependence between two signals in the frequency domain; and (3) Phase-locking value (PLV) assesses the consistency of instantaneous phase differences independent of signal amplitude.

**EEG signals**

EEG data were preprocessed using EEGLAB toolbox[5]. Following the removal of non-scalp channels and the rejection of bad segments, signals were bandpass filtered (0.5–45 Hz) using a standard finite impulse response (FIR) filter with zero-phase delay. The data were re-referenced to the Cz electrode, downsampled to 250 Hz for computational efficiency. Independent component analysis (ICA) was applied to identify and eliminate ocular, muscular, and cardiac artifacts and then z-score normalized for each channel.

**Individualized FEM-based forward modeling**

Individualized EEG forward modeling used SimNIBS 4.1.0 to generate the realistic volume conductor model[6]. High-quality individualized head meshes were reconstructed from the T1w of each participant, segmenting the head into multiple tissue compartments (white matter, gray matter, CSF, skull, and scalp) with distinct conductivities. By solving the potential distributions across tetrahedral elements of meshes, we computed individualized leadfield matrices. For the standardized control model, the identical pipeline was applied to the MNI152 standard template.

**Signal complexity**

To characterize the dynamical complexity of the EEG signals, three complementary non-linear time-series measures were computed channel-wise, including Lempel–Ziv complexity (LZC), the Hurst exponent, and permutation–fuzzy entropy (PFEn). For LZC, signals were mean-centered, median-binarized, and evaluated using the exhaustive parsing algorithm. Long-range temporal dependence was quantified via the Hurst exponent, estimated using rescaled range analysis across 20 logarithmically spaced scales. Local irregularity was assessed using PFEn, defined as the window-averaged sum of fuzzy and permutation entropy computed over non-overlapping windows.

**Power spectral density**

The power spectral density (PSD) of the EEG signals (resampled to 250 Hz) was estimated utilizing Welch's method. Signals were segmented into 2-second epochs with 50% overlap and tapered with a Hamming window to minimize spectral leakage. The resulting PSDs were averaged across epochs within the 0.5–45 Hz range for both empirical and simulated data.

**EEG microstates**

EEG microstates were extracted to analyze large-scale spatiotemporal dynamics[7]. Following standard preprocessing, aggregated empirical global field power peaks were clustered into four canonical templates using modified K-means. These empirical templates serve solely as a fixed spatial coordinate system. Individual continuous EEG data were then back-fitted to these templates (with 30-ms smoothing) to compute standard microstate characteristics (global explained variance, coverage, occurrence, duration, and transition probabilities). For comparison, the identically preprocessed simulated EEG signals were directly back-projected onto the empirically derived templates. It is critical to note that while empirical and simulated continuous signals share these spatial templates for evaluation, all resulting temporal dynamics (e.g., transition probabilities, occurrence, and duration) were strictly held out from the model optimization, ensuring an independent validation of the emergent spatiotemporal behavior.

### Supplementary figures and tables

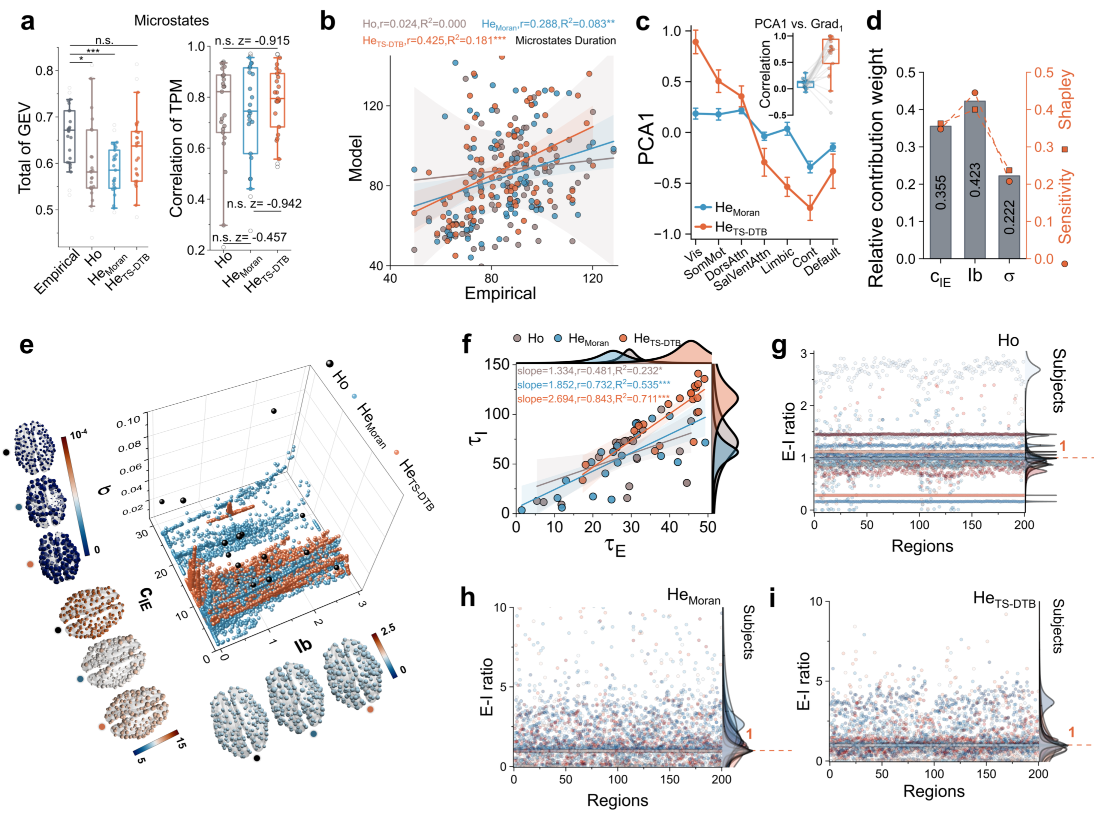


**Fig. S1. Comparative analysis of homogeneous and heterogeneous models. (a)** EEG microstate metrics (global explained variance and transition probability matrix correlation). **(b)** Microstate duration fitting results. **(c)** Alignment of parameter space topology with the empirical functional hierarchy in the TS-DTB models (orange) versus null models (blue). **(d)** Parameter contribution analysis for the E-I ratio, including relative weights, sensitivity, and Shapley values. **(e)** Three-dimensional individualized parameter landscapes and group-level brain topographies under gradient priors. **(f)** Time constant distributions and linear fits across models. **(g–i)** Regional E-I ratio distributions across individuals under the three modeling schemes. In (a, c), boxplots show the median and IQR across n = 25 biologically independent HC individuals (two-sided paired t-tests, $*p < 0.05$, $***p < 0.001$; n.s., not significant). In (b, f), solid lines indicate linear regression fits with 95% confidence intervals (shaded).

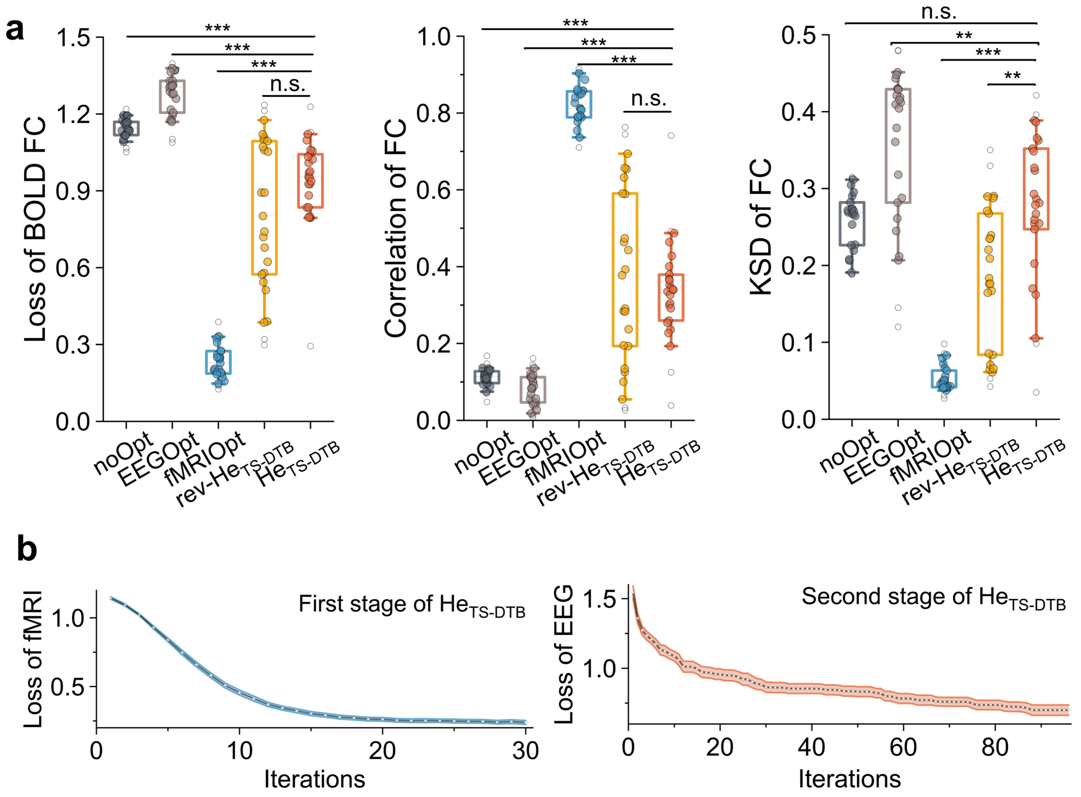


**Fig. S2. Detailed breakdown of fMRI functional connectivity fitting metrics and optimization convergence curves. (a)** Quantitative evaluation of the total loss of BOLD FC, correlation of FC, and KSD of FC across five parameterization schemes: the unconstrained baseline (noOpt), the EEG-constrained models (EEGOpt), the first-stage fMRI-constrained models (fMRIOpt), the final proposed models (He$_{TS\text{-}DTB}$), and the reversed-order optimization models (rev-He$_{TS\text{-}DTB}$). Boxplots indicate the median (center line) and interquartile range (box limits). Whiskers represent the 90% confidence intervals, and dots denote biologically independent individuals (n = 25). Significance was assessed via two-sided paired t-tests (*$p$ < 0.05, **$p$ < 0.01, ***$p$ < 0.001, n.s. indicates not significant). **(b)** Convergence curves for the first-stage iterative fMRI updates (left) and second-stage Bayesian EEG optimization (right). Solid lines represent the group mean loss, and shaded areas indicate the SE across subjects (n = 25).

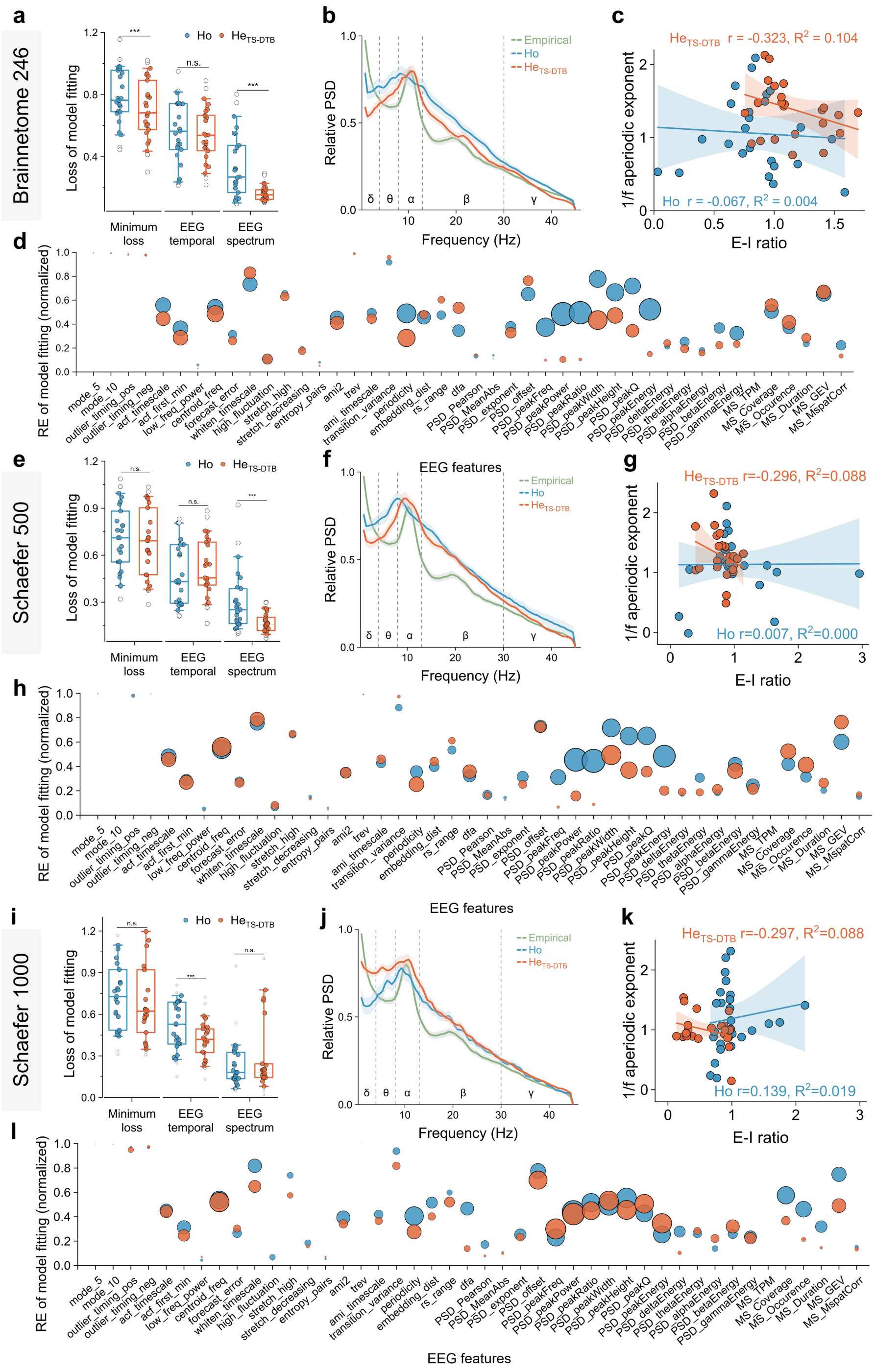


**Fig. S3. Cross-parcellation stability of the TS-DTB models.** Evaluation of optimization performance and mechanistic fidelity across the Brainnetome 246, Schaefer 500, and Schaefer 1000 atlases. **(a, e, i)** Comparison of minimum loss and median RE between homogeneous (Ho) and heterogeneous ($He_{TS\text{-}DTB}$) models. Boxplots indicate the median and IQR (n = 25 biologically independent HC individuals; two-sided paired t-tests, **p* < 0.01, ****p* < 0.001; n.s., not significant). **(b, f, j)** Channel-averaged relative PSD for empirical data and simulations (solid curves represent the group mean and shaded areas indicate the SE, n = 25). **(c, g, k)** Scatter plots showing the negative correlation between the simulated E-I ratio and the aperiodic exponent across all three atlases (solid lines indicate linear fits with 95% confidence intervals, n = 25). (d, h, l) Feature-wise REs (circle centers indicate the mean RE and diameters are proportional to the SE).

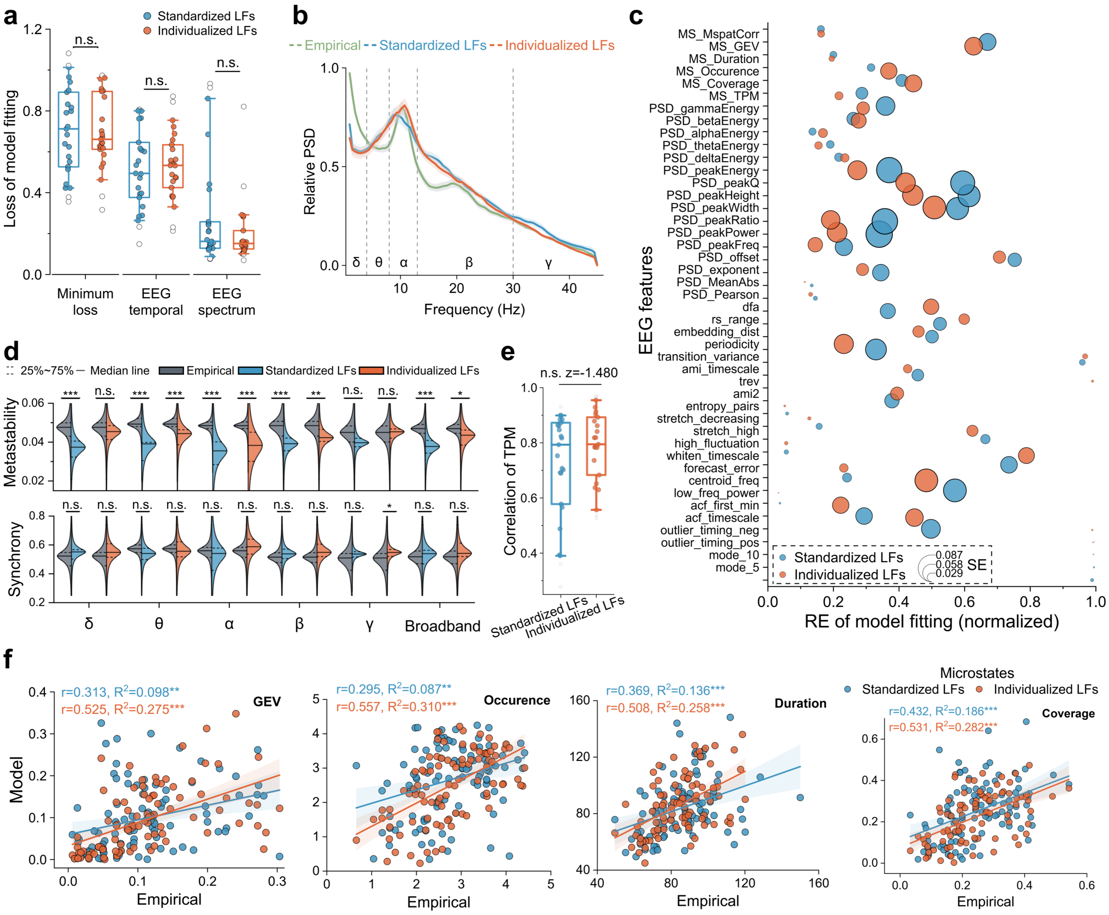


**Fig. S4. Impact of individualized versus standardized EEG forward modeling on simulation fidelity.** Quantitative comparison using leadfields derived from subject-specific MRI (individualized LFs) versus a template MNI head model (standardized LFs) across n = 25 biologically independent HC individuals. **(a)** Optimization performance (minimum loss and median RE). **(b)** Channel-averaged relative PSD (mean ± SE). **(c)** Feature-wise REs for EEG temporal, spectral, and microstate properties (circle centers and diameters denote mean and SE). **(d)** Violin plots showing the distribution of metastability (top) and synchrony (bottom) across frequency bands for empirical data (grey) and simulations (blue/orange). **(e)** Correlation of the microstate transition probability matrix (TPM) between empirical and simulated data. **(f)** Scatter plots relating empirical and simulated microstate metrics (GEV, occurrence, duration, and coverage), with linear regression fits and 95% confidence intervals (shaded). In (a, d, e), boxplots/violins show the median and IQR, with significance assessed via two-sided paired t-tests (*$p$ < 0.05, ***$p$ < 0.001; n.s., not significant).

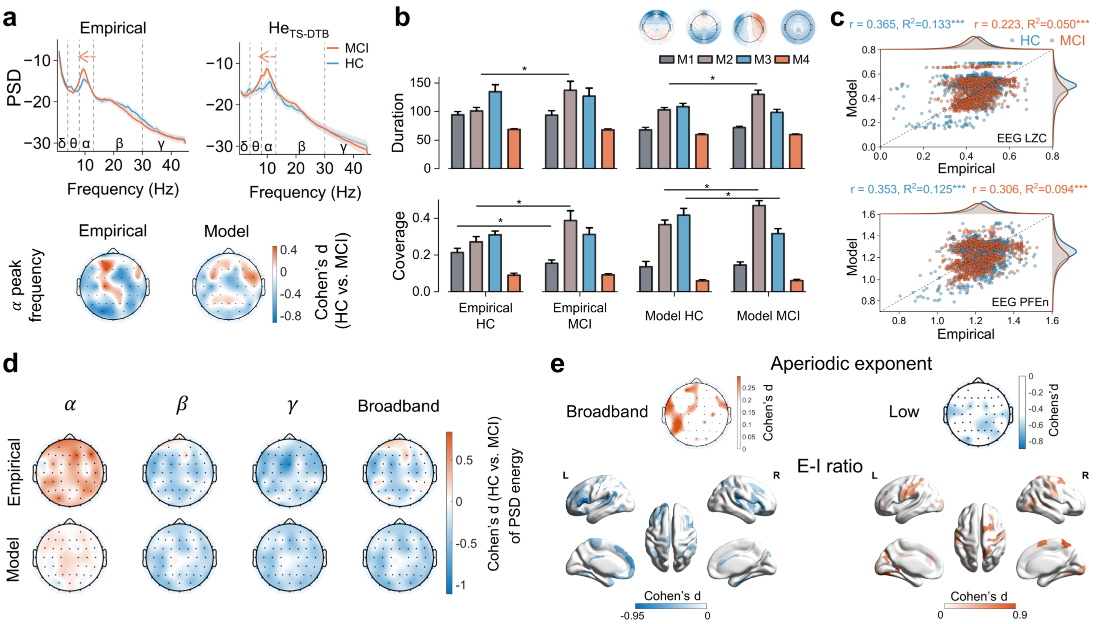


**Fig. S5. Validation in the mild cognitive impairment (MCI) cohort. (a)** Channel-averaged PSD (mean ± SE) for empirical data and $He_{TS\text{-}DTB}$ models (top), alongside Cohen's d maps for alpha peak frequency (bottom). **(b)** Microstate duration and coverage metrics (bars and error bars indicate mean ± SE across n = 16 HC and n = 15 MCI biologically independent individuals; two-sided independent samples t-tests, $*p < 0.05$). **(c)** Scatter plots correlating empirical and simulated LZC and permutation-fuzzy entropy (PFEn) (linear fits with 95% confidence intervals). **(d)** Cohen's d maps of spectral energy alterations across frequency bands. (e) Spatial correspondence between residual aperiodic exponent effects (low: delta to alpha band) and local E-I ratios.

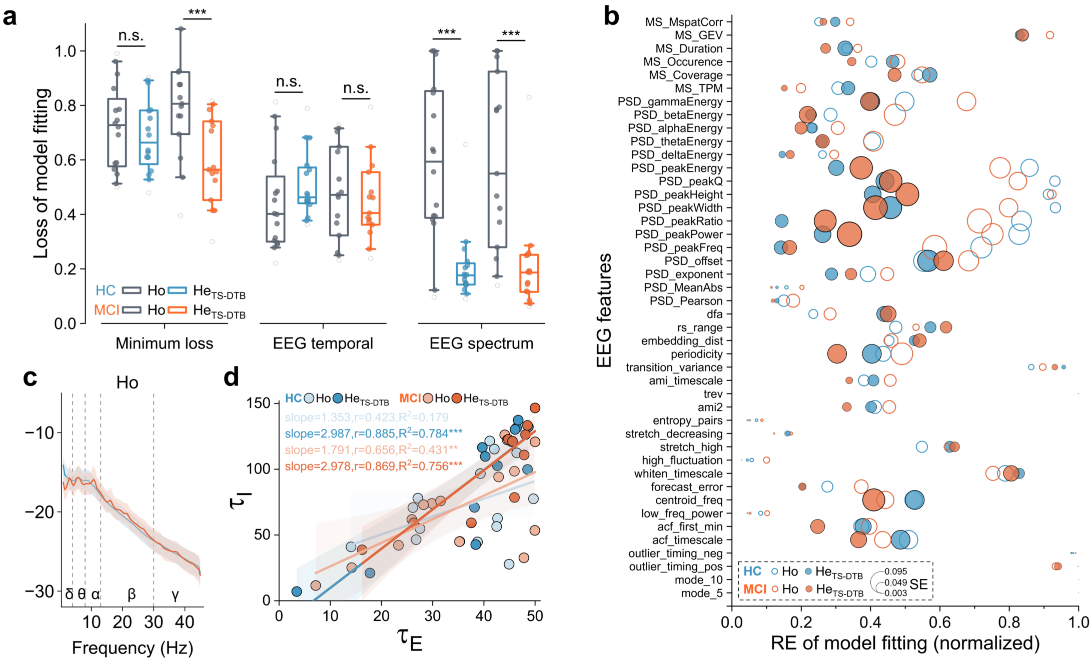


**Fig. S6. Cross-dataset stability of the TS-DTB in the MCI cohort. (a)** Optimization performance (loss and REs) across parameterization schemes and groups. Boxplots indicate the median and IQR across n = 16 HC and n = 15 MCI biologically independent individuals (two-sided paired t-tests, ***$p$ < 0.001; n.s., not significant). **(b)** Feature-wise REs for EEG temporal, spectral, and microstate properties (circle centers and diameters indicate mean and SE). **(c)** Channel-averaged PSD (mean ± SE) for homogeneous (Ho) models. **(d)** Distributions and linear fits (with 95% confidence intervals) of time constants in HC and MCI participants.

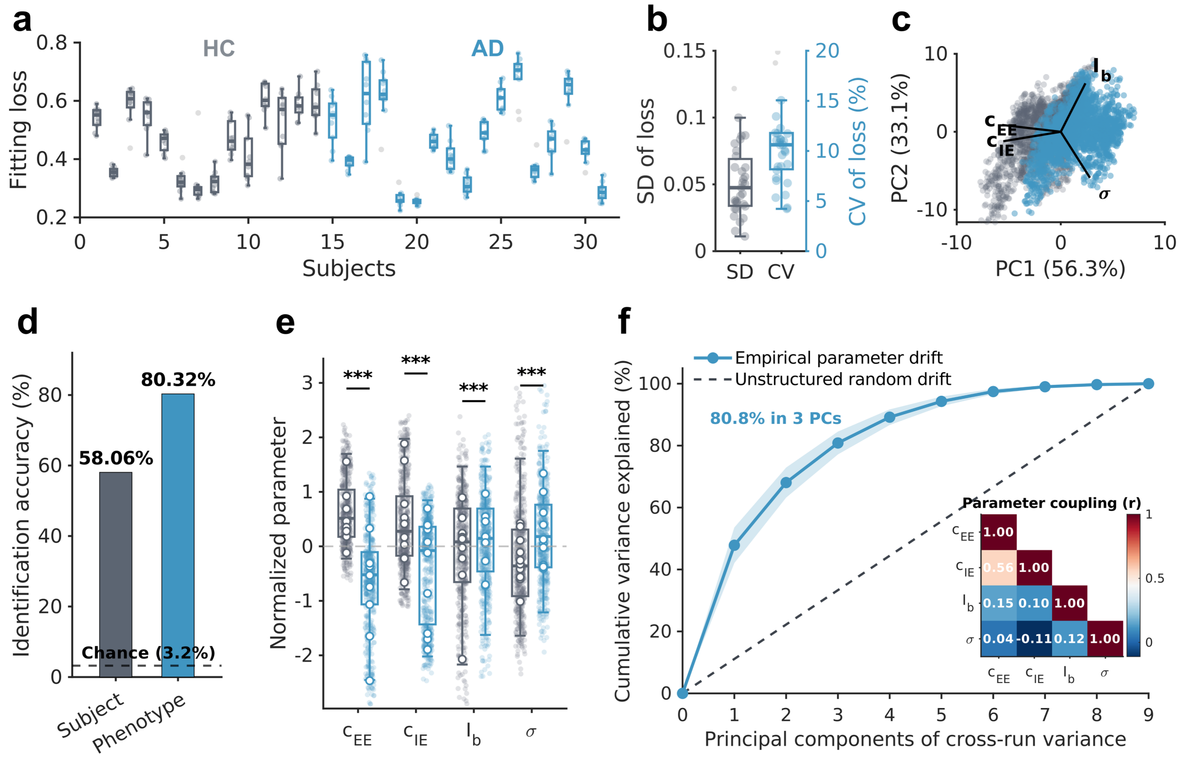


**Fig. S7. Parameter analysis across 10 independent Bayesian optimization runs. (a)** Overall loss function values across 10 independent random initializations for each subject (n = 14 HC and n = 17 AD). Boxplots indicate the median (center line) and IQR (box limits), whiskers show 1.5 × IQR, and dots denote individual runs. **(b)** Distribution of the Standard Deviation (SD) and Coefficient of Variation (CV) of the loss across the 10 runs for all subjects (n = 31). **(c)** PCA projection of the 800-dimensional regional parameters across all runs (n = 310), maintaining spatial separation between HC (gray) and AD (blue) phenotypes. **(d)** Nearest-neighbor identification accuracy for individual subjects (58.06%) and clinical phenotypes (80.32%) compared against the chance level (3.2%, dashed line). **(e)** Standardized distributions of the four biophysical parameters across 10 runs, with transparent lines connecting matched-seed group means (two-sided paired samples t-tests, $^{***}p < 0.001$). **(f)** Cumulative variance explained by the principal components of cross-run parameter drift (blue line, mean ± SE) versus random noise (dashed line), with 80.8% of variance restricted to 3 compensatory axes. Inset: cross-seed parameter coupling matrix (Pearson's $r$).

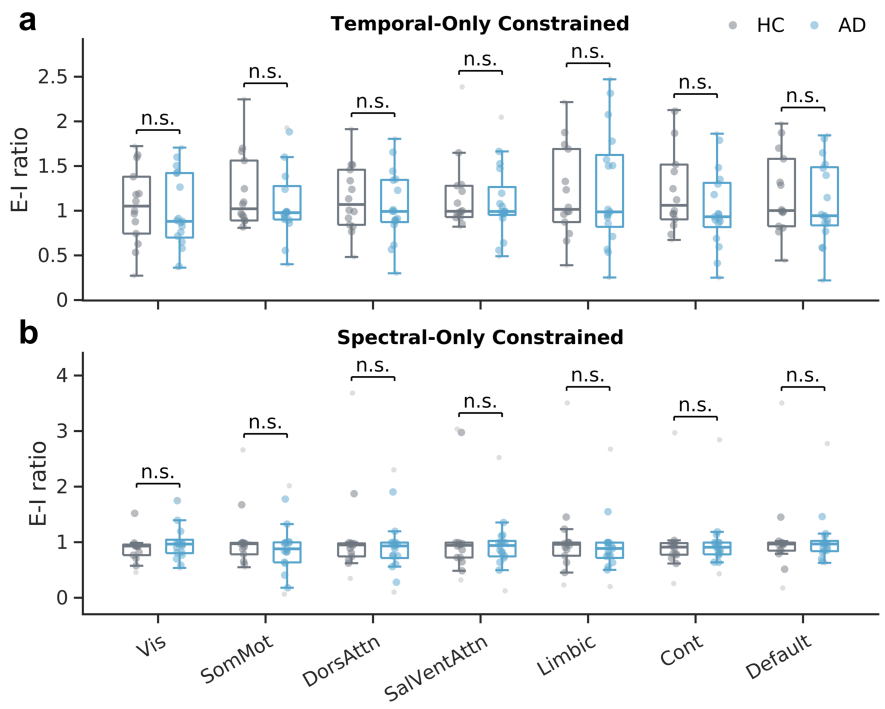


**Fig. S8. Feature hold-out validation for local E-I ratios under single-domain constraints.** Individualized models were independently re-optimized using **(a)** Temporal-Only constraints (holding out PSD features) and **(b)** Spectral-Only constraints (holding out catch22 features). Single-domain optimizations failed to identify significant E-I ratio differences between HC and AD across functional subnetworks. Boxplots indicate the median (center line) and IQR (box limits), whiskers represent 1.5 × IQR, and dots denote biologically independent individuals (n = 14 HC, n = 17 AD). Significance was assessed via two-sided independent samples t-tests (n.s. indicates not significant).

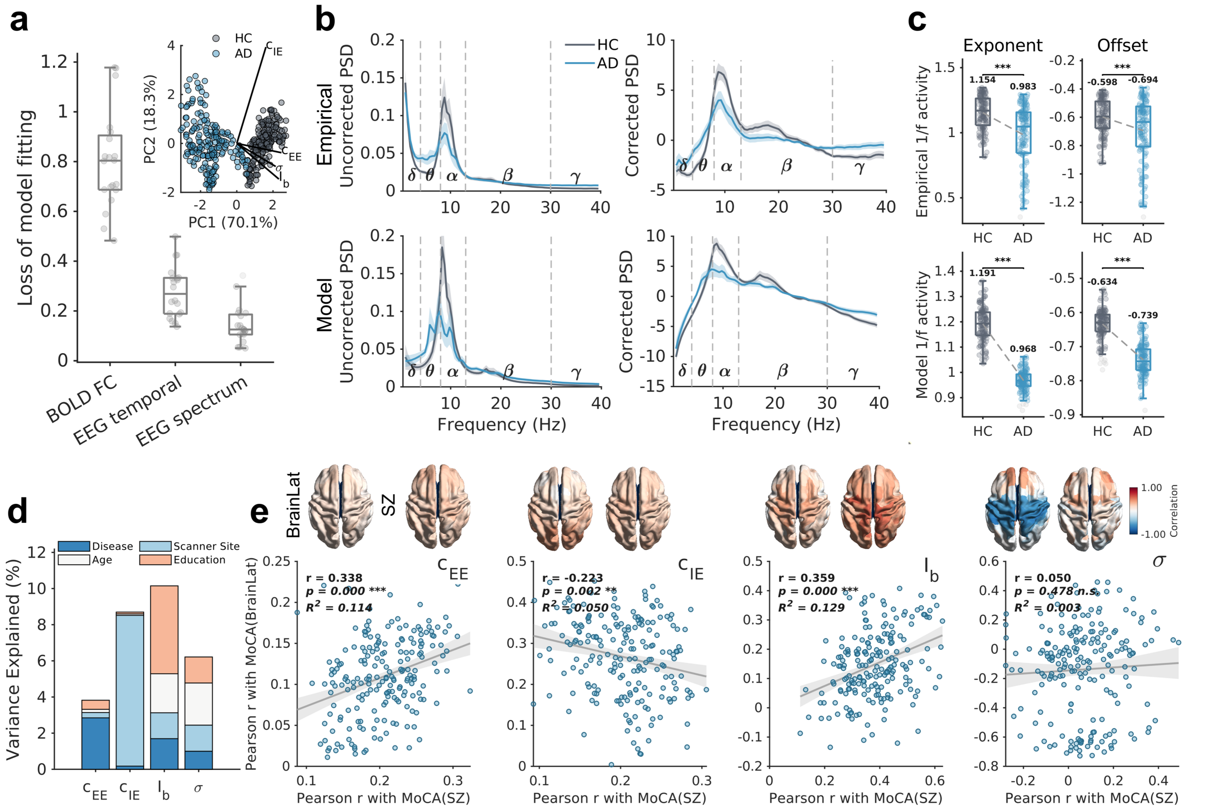


**Fig. S9. Cross-site external validation and parameter assessment in the BrainLat cohort** (n = 10 HC, n = 15 AD). **(a)** Loss function values for BOLD FC, EEG temporal, and EEG spectral fitting stages. Inset: PCA projection of optimized regional parameters with parameter loading vectors (black lines). **(b)** Empirical (top) and simulated (bottom) uncorrected (left) and aperiodic-corrected (right) PSD profiles. Solid curves and shaded areas represent the group mean ± SEM. **(c)** Empirical (top) and simulated (bottom) aperiodic exponent and offset distributions. **(d)** Region-wise Variance Partitioning Analysis (VPA) averaged across cortical regions, quantifying the independent parameter variance explained by Disease, Scanner Site, Age, and Education. **(e)** Cross-site spatial consistency of regional parameter–MoCA correlations between the SZ and BrainLat cohorts across n = 200 cortical regions (solid lines and shaded bands indicate linear fits and 95% confidence intervals). For all boxplots, the center line indicates the median, box limits represent the IQR, and whiskers extend to 1.5 × IQR. Significance in (c) was assessed via channel-wise two-sided paired t-tests (n = 128 channels; $**p < 0.01$, $***p < 0.001$, n.s., not significant).

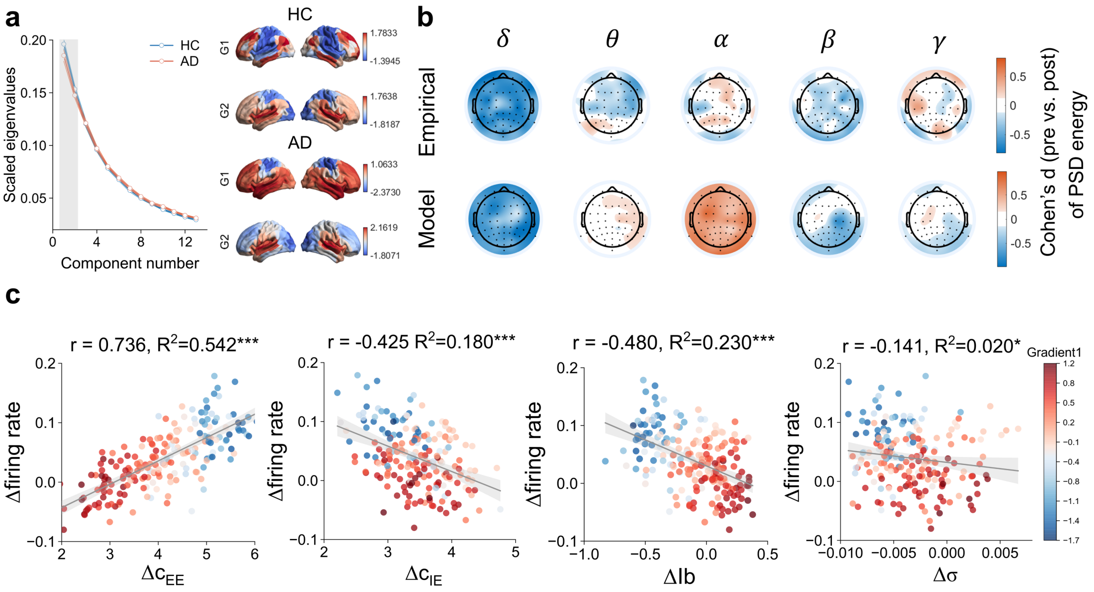


**Fig. S10. Results of rTMS regulation studies in the AD cohort. (a)** Scaled eigenvalues as a function of the gradient component number (blue, HC; orange, AD; mean ± SE across n = 14 HC and n = 17 AD individuals). **(b)** Topographic Cohen's d maps of PSD energy changes post-rTMS for empirical data (top) and model simulations (bottom). **(c)** Scatter plots correlating model parameter shifts ($\Delta c_{EE}$, $\Delta c_{IE}$, $\Delta Ib$ and $\Delta\sigma$) with firing rate changes across n = 200 cortical regions (solid lines indicate linear regression fits with 95% confidence intervals; $*p < 0.05$, $***p < 0.001$). Points are colored by the first functional gradient (blue, low gradient; red, high gradient).

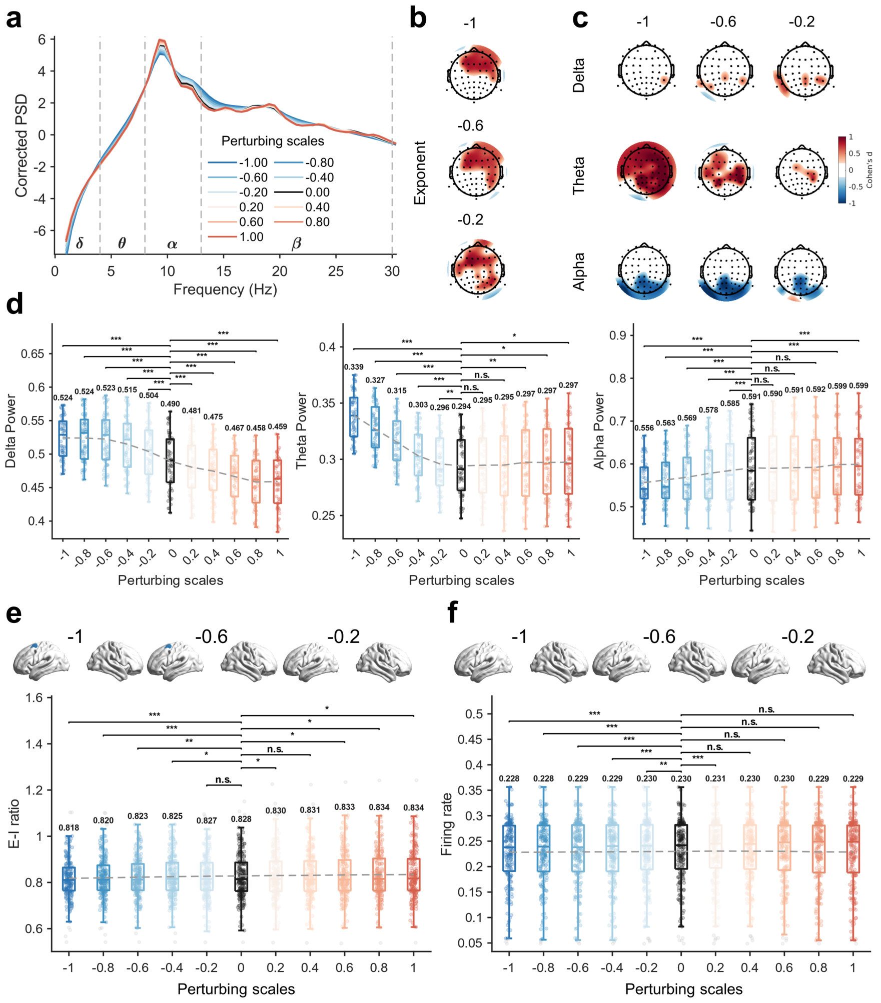


**Fig. S11. Parameter perturbation in healthy control models.** Individualized healthy control (HC) models (n = 14) were perturbed along the empirical group-difference vector ($\Delta\sigma_{HC-AD}$) using scaling factors $s \in [-1.0,1.0]$ ($\sigma_{HC-AD} = \sigma_{HC} + s \cdot \Delta\sigma_{HC-AD}$), where $s = 0$ denotes the unperturbed HC baseline and $s < 0$ shifts the model toward the AD regime. **(a)** Channel-averaged periodic PSD (Corrected PSD) across perturbation scales (solid curves represent the group mean). **(b, c)** Topographic Cohen's d maps comparing negative scales ($s \in$ [-1.0, -0.2]) against baseline ($s = 0$) for the aperiodic exponent (b) and delta, theta, and alpha power (c). **(d)** Channel-wise power distributions in the delta, theta, and alpha bands across perturbation scales. **(e, f)** Region-wise distributions of the model-inferred E-I ratio (e) and simulated excitatory firing rates (f) across perturbation scales, with corresponding cortical surface Cohen's d maps for negative scales (= -1.0, -0.6, -0.2 vs. baseline $s = 0$). For all boxplots in (d–f), the center line indicates the median, box limits represent the IQR, whiskers extend to 1.5 × IQR, numerical labels indicate group means, and dots denote EEG channels in (d) (n = 61 channels) or cortical regions in (e, f) (n = 200 regions) averaged across n = 14 biologically independent HC subjects. Significance relative to baseline ($s = 0$) was assessed via two-sided paired t-tests (*$p < 0.05$, **$p < 0.01$, ***$p < 0.001$; n.s., not significant).

#### Parameter sets and feature sets

The detailed configuration of the model optimization and the multidimensional feature sets employed to evaluate model performance are shown in Tables S1-S4. To ensure biological plausibility and computational tractability, we defined specific search ranges and prior distributions for the key biophysical parameters governing local circuit dynamics (Table S1). Crucially, we maintained a strict separation between features used for model fitting and those reserved for independent validation. The loss function used to constrain the second-stage model optimization (Eq. 14 in the main text) was constructed exclusively from two local spectral-temporal feature sets: the catch22 time-series feature set capturing linear and non-linear temporal properties (Table S2) and a spectral feature set characterizing the power spectral density (PSD) morphology, including periodic and aperiodic components (Table S3). In contrast, the EEG microstate feature set (Table S4), which quantifies the global spatiotemporal syntax of large-scale brain networks, was strictly held out from the parameter optimization process and served exclusively as an independent out-of-sample validation of the model's emergent macroscopic dynamics.

**Table S1. Parameter descriptions for Bayesian optimization**[8]

| Parameter | Symbol | Prior | Nominal |
|---|---|---|---|
| Excitatory time constant | $\tau_E$ | U (1, 50) | 8 ms |
| Ratio factor of the inhibitory time constant | $r_{\tau_I}$ | U (0.5, 3) | 8 ms ($\tau_I$) |
| Global coupling strength | $G_e$ | U (0.5, 1.5) | 1.0 |
| Inhibitory to excitatory synaptic strength | $c_{IE}$ | U (3, 36) | 12 |
| Background input to populations | $Ib$ | U (0, 3) | 1.15 |
| Exponent of Gaussian noise standard deviation | $\sigma$ | U (-4, -1) | -1 |
| Linear coefficients of $c_{IE}$ | $a_k^{cIE}$ | U (-20, 20) | |
| Linear coefficients of $c_{IE}$ | $b_k^{cIE}$ | U (-20, 20) | |
| Linear coefficients of $c_{IE}$ | $c_k^{cIE}$ | U (3, 30) | |
| Linear coefficients of $Ib$ | $a_k^{Ib}$ | U (-20, 20) | |
| Linear coefficients of $Ib$ | $b_k^{Ib}$ | U (-20, 20) | |
| Linear coefficients of $Ib$ | $c_k^{Ib}$ | U (0, 2) | |
| Linear coefficients of $\sigma$ | $a_k^{\sigma}$ | U (-5, 5) | |
| Linear coefficients of $\sigma$ | $b_k^{\sigma}$ | U (-5, 5) | |
| Linear coefficients of $\sigma$ | $c_k^{\sigma}$ | U (-3, -2) | |
| Linear coefficients of $c_{EE}$ | $a_k^{cEE}$ | U (-20, 20) | |
| Linear coefficients of $c_{EE}$ | $b_k^{cEE}$ | U (-20, 20) | |
| Linear coefficients of $c_{EE}$ | $c_k^{cEE}$ | U (1, 48) | |

**Table S2. The overview of the catch22 temporal feature sets[9]**

| | Feature short name | Category | Description |
|---|---|---|---|
| 1 | mode_5 | Distribution shape | 5-bin histogram mode |
| 2 | mode_10 | Distribution shape | 10-bin histogram mode |
| 3 | outlier_timing_pos | Extreme event timing | Positive outlier timing |
| 4 | outlier_timing_neg | Extreme event timing | Negative outlier timing |
| 5 | acf_timescale | Linear autocorrelation | First $1/e$ crossing of the ACF |
| 6 | acf_first_min | Linear autocorrelation | First minimum of the ACF |
| 7 | low_freq_power | Linear autocorrelation | Power in lowest 20% frequencies |
| 8 | centroid_freq | Linear autocorrelation | Centroid frequency |
| 9 | forecast_error | Simple forecasting | Error of 3-point rolling mean forecast |
| 10 | whiten_timescale | Incremental differences | Change in autocorrelation timescale after incremental differencing |
| 11 | high_fluctuation | Incremental differences | Proportion of high incremental changes in the series |
| 12 | stretch_high | Symbolic | Longest stretch of above-mean values |
| 13 | stretch_decreasing | Symbolic | Longest stretch of decreasing values |
| 14 | entropy_pairs | Symbolic | Entropy of successive pairs in symbolized series |
| 15 | ami2 | Nonlinear autocorrelation | Histogram-based automutual information (lag 2, 5 bins) |
| 16 | trev | Nonlinear autocorrelation | Time reversibility |
| 17 | ami_timescale | Linear autocorrelation structure | First minimum of the AMI function |
| 18 | transition_variance | Symbolic | Transition matrix column variance |
| 19 | periodicity | Linear autocorrelation structure | Wang's periodicity metric |
| 20 | embedding_dist | Other | Goodness of exponential fit to embedding distance distribution |
| 21 | rs_range | Self-affine scaling | Rescaled range fluctuation analysis (low-scale scaling) |
| 22 | dfa | Self-affine scaling | Detrended fluctuation analysis (low-scale scaling) |

**Table S3. The overview of spectrum feature sets[10]**

| | Feature short name | Category | Description |
|---|---|---|---|
| 1 | PSD_Pearson | Broadband similarity | Pearson correlation between empirical and simulated PSD |
| 2 | PSD_MeanAbs | Broadband deviation | Mean absolute deviation between empirical and simulated PSD |
| 3 | PSD_exponent | Aperiodic component | Exponent of the aperiodic $1/f$ derived from spectral parameterization |
| 4 | PSD_offset | Aperiodic component | Vertical offset of the aperiodic $1/f$ derived from spectral parameterization |
| 5 | PSD_peakFreq | Oscillatory peak | Center frequency of the dominant oscillatory peak |
| 6 | PSD_peakPower | Oscillatory peak | Absolute power at the dominant peak frequency |
| 7 | PSD_peakRatio | Oscillatory peak | Ratio of peak power to aperiodic background power |
| 8 | PSD_peakWidth | Oscillatory peak | Full width at half maximum (FWHM) of the dominant peak |
| 9 | PSD_peakHeight | Oscillatory peak | Prominence of the dominant peak |
| 10 | PSD_peakQ | Oscillatory peak | Peak quality factor ($f_{peak}/peak_{width}$) |
| 11 | PSD_peakEnergy | Oscillatory peak | Integrated spectral energy within the peak-centered window (±1 Hz) |
| 12 | PSD_deltaEnergy | Band-limited energy | Spectral energy in the delta band (0.5–4 Hz) |
| 13 | PSD_thetaEnergy | Band-limited energy | Spectral energy in the theta band (4–8 Hz) |
| 14 | PSD_alphaEnergy | Band-limited energy | Spectral energy in the alpha band (8–13 Hz) |
| 15 | PSD_betaEnergy | Band-limited energy | Spectral energy in the beta band (13–30 Hz) |
| 16 | PSD_gammaEnergy | Band-limited energy | Spectral energy in the gamma band (30–45 Hz) |

**Table S4. The overview of EEG microstate feature sets[7]**

| | Feature short name | Category | Description |
|---|---|---|---|
| 1 | MS_TPM | Temporal dynamics | Transition probabilities between microstate classes |
| 2 | MS_Coverage | Temporal stability | Proportion of total time spent in each class |
| 3 | MS_Occurrence | Temporal dynamics | Number of state entries per second |
| 4 | MS_Duration | Temporal stability | Mean duration of each state segment |
| 5 | MS_GEV | Explanatory power | Variance of EEG topographies explained by each class |
| 6 | MS_MspatCor | Spatial similarity | Spatial correlation between templates and assigned maps |

**Table S5. Demographic and clinical profile of healthy controls in the AD cohort**

| ID | Gender | Age | Education | MMSE | HAMA | HAMD |
|---|---|---|---|---|---|---|
| 1 | Female | 50 | 12 | 28 | 3 | 2 |
| 2 | Female | 50 | 6 | 29 | 4 | 1 |
| 3 | Female | 52 | 9 | 30 | 0 | 0 |
| 4 | Female | 56 | 16 | 30 | 0 | 0 |
| 5 | Female | 56 | 9 | 29 | 0 | 0 |
| 6 | Male | 61 | 15 | 29 | 0 | 0 |
| 7 | Female | 56 | 0 | 27 | 12 | 11 |
| 8 | Female | 62 | 15 | 30 | 5 | 0 |
| 9 | Female | 56 | 15 | 30 | 2 | 1 |
| 10 | Female | 63 | 9 | 30 | 4 | 4 |
| 11 | Female | 67 | 9 | 28 | 8 | 4 |
| 12 | Female | 57 | 12 | 28 | 4 | 6 |
| 13 | Female | 53 | 9 | 30 | 2 | 0 |
| 14 | Male | 50 | 15 | 30 | 2 | 0 |
| | 2/12(Male/Female) | 56.36±5.29 | 10.79±4.44 | 29.14±1.03 | 3.29±3.41 | 2.07±3.22 |

**Table S6. Demographic and clinical profile of AD in the AD cohort**

| ID | Gender | Age | Education | MOCA | MMSE | HAMA | HAMD | Years of illness |
|---|---|---|---|---|---|---|---|---|
| 1 | Female | 46 | 16 | 30 | 29 | 7 | 4 | 2 |
| 2 | Female | 65 | 0 | 10 | 19 | 2 | 2 | 0.8 |
| 3 | Female | 64 | 12 | 22 | 26 | 4 | 8 | 1 |
| 4 | Male | 73 | 20 | 16 | 27 | 5 | 6 | 1 |
| 5 | Female | 66 | 12 | 17 | 25 | 10 | 10 | 2 |
| 6 | Male | 68 | 12 | 23 | 26 | 11 | 9 | 2 |
| 7 | Female | 73 | 12 | 26 | 30 | 10 | 14 | 2 |
| 8 | Female | 50 | 9 | 23 | 30 | 21 | 18 | 1 |
| 9 | Female | 57 | 9 | 23 | 29 | 9 | 9 | 2.5 |
| 10 | Female | 71 | 0 | 10 | 17 | 10 | 14 | 3 |
| 11 | Female | 71 | 6 | 18 | 25 | 3 | 7 | 4 |
| 12 | Female | 67 | 6 | 18 | 25 | 6 | 2 | 2 |
| 13 | Female | 83 | 6 | 7 | 13 | 7 | 7 | 2 |
| 14 | Female | 53 | 20 | 27 | 29 | 4 | 7 | 1 |
| 15 | Female | 75 | 9 | 12 | 17 | 4 | 3 | 3.5 |
| 16 | Male | 65 | 9 | 13 | 24 | 0 | 2 | 1 |
| 17 | Female | 67 | 0 | 25 | 21 | 11 | 8 | 0.5 |
| | 3/14(Male/Female) | 65.53±9.46 | 9.88±5.80 | 18.8±6.77 | 24.24±5.12 | 7.29±4.87 | 7.65±4.55 | 1.84±0.99 |

**Table S7. Demographic and clinical profile of healthy controls in the MCI cohort**

| ID | Gender | Age | Education | MoCA-B | ADAS-cog |
|---|---|---|---|---|---|
| 1 | Female | 65 | 13 | 29 | 7.333 |
| 2 | Female | 59 | 17 | 28 | 4.330 |
| 3 | Female | 57 | 12 | 24 | 4.667 |
| 4 | Male | 69 | 7 | 23 | 8.300 |
| 5 | Female | 67 | 9 | 28 | 11.300 |
| 6 | Male | 63 | 15 | 27 | 4.000 |
| 7 | Female | 59 | 14 | 27 | 5.667 |
| 8 | Male | 60 | 13 | 25 | 3.667 |
| 9 | Female | 70 | 12 | 26 | 6.333 |
| 10 | Female | 70 | 18 | 29 | 2.333 |
| 11 | Female | 51 | 15 | 28 | 2.333 |
| 12 | Female | 63 | 18 | 28 | 1.667 |
| 13 | Female | 69 | 13 | 28 | 4.333 |
| 14 | Female | 64 | 16 | 28 | 6.667 |
| 15 | Female | 58 | 14 | 26 | 6.000 |
| 16 | Male | 58 | 10 | 25 | 5.333 |
| | 4/12(Male/Female) | 62.63±5.54 | 13.50±3.10 | 26.81±1.80 | 5.27±2.45 |

**Table S8. Demographic and clinical profile of MCI in the MCI cohort**

| ID | Gender | Age | Education | MoCA-B | ADAS-cog | ADAS-10 |
|---|---|---|---|---|---|---|
| 1 | Female | 59 | 12 | 26 | 8.000 | 5.333 |
| 2 | Male | 68 | 15 | 21 | 7.367 | 2.700 |
| 3 | Female | 58 | 7 | 21 | 13.333 | 4.667 |
| 4 | Male | 65 | 12 | 23 | 7.333 | 4.333 |
| 5 | Female | 70 | 11 | 25 | 12.333 | 5.000 |
| 6 | Female | 70 | 13 | 27 | 12.000 | 5.000 |
| 7 | Female | 68 | 7 | 22 | 13.667 | 4.333 |
| 8 | Male | 66 | 12 | 18 | 12.000 | 6.667 |
| 9 | Male | 54 | 15 | 24 | 7.333 | 4.667 |
| 10 | Female | 62 | 16 | 25 | 8.333 | 4.333 |
| 11 | Female | 59 | 9 | 20 | 11.000 | 2.667 |
| 12 | Female | 65 | 15 | 21 | 20.667 | 5.000 |
| 13 | Male | 71 | 12 | 25 | 9.333 | 4.000 |
| 14 | Male | 69 | 15 | 22 | 6.000 | 5.333 |
| 15 | Male | 67 | 9 | 25 | 11.667 | 3.667 |
| | 7/8(Male/Female) | 64.73±5.18 | 12.00±2.95 | 23.00±2.54 | 10.69±3.71 | 4.51±1.02 |

**Table S9. Demographic and clinical profile of healthy controls in the BrainLat cohort**

| ID | Gender | Age | Education | MoCA |
|---|---|---|---|---|
| 1 | Female | 72 | 12 | 24 |
| 2 | Male | 76 | 10 | 17 |
| 3 | Male | 70 | 16 | 29 |
| 4 | Female | 80 | 10 | 24 |
| 5 | Male | 72 | 17 | 22 |
| 6 | Female | 73 | 11 | 23 |
| 7 | Male | 73 | 13 | 23 |
| 8 | Male | 74 | 8 | 24 |
| 9 | Female | 77 | 17 | 27 |
| 10 | Female | 77 | 22 | 29 |
| | 5/5(Male/Female) | 74.40±3.03 | 13.60±4.30 | 24.22±3.77 |

Note: Missing MoCA values for subject 1 were replaced with the group average.

**Table S10. Demographic and clinical profile of AD in the BrainLat cohort**

| ID | Gender | Age | Education | MoCA |
|---|---|---|---|---|
| 1 | Male | 76 | 12 | 10 |
| 2 | Male | 71 | 15 | 17 |
| 3 | Female | 81 | 11 | 17 |
| 4 | Male | 76 | 5 | 17 |
| 5 | Female | 82 | 12 | 15 |
| 6 | Male | 86 | 12 | 14 |
| 7 | Female | 98 | 16 | 18 |
| 8 | Female | 79 | 8 | 16 |
| 9 | Female | 76 | 12 | 18 |
| 10 | Female | 80 | 9 | 20 |
| 11 | Male | 70 | 4 | 15 |
| 12 | Female | 69 | 12 | 19 |
| 13 | Female | 66 | 8 | 15 |
| 14 | Female | 90 | 6 | 17 |
| 15 | Female | 64 | 16 | 21 |
| | 5/10(Male/Female) | 77.60±9.19 | 10.53±3.78 | 16.54±2.88 |

Note: Missing MoCA values for subjects 3 and 4 were replaced with the group average.
